# The essential roles of Dicer-mediated mTORC1 signaling in parathyroid gland integrity and function: Insights from genetic mouse models and clinical data

**DOI:** 10.1101/2023.08.21.554016

**Authors:** Alia Hassan, Nareman Khalaily, Rachel Kilav-Levin, Barbara Del Castello, Nancy Ruth Manley, Iddo Z. Ben-Dov, Tally Naveh-Many

## Abstract

Secondary hyperparathyroidism (SHP) frequently accompanies chronic kidney disease (CKD), contributing to morbidity and mortality in patients. Our previous findings demonstrated that PT-*Dicer^-/-^* mice, with parathyroid specific deletion of *Dicer* and consequently microRNA, maintained normal serum PTH levels but failed to increase serum PTH in response to the major inducers of PTH secretion, hypocalcemia and CKD. Additionally, we elucidated a critical role of mTORC1 in CKD-induced SHP. We now explored the roles of Dicer and mTORC1 in parathyroid development and function. Despite sustaining normal serum PTH levels, PT-*Dicer^-/-^* mice displayed apoptotic loss of intact parathyroid glands postnatally, which were replaced by scattered cell clusters, and reduced mTORC1 activity. PT-*mTORC1^-/-^* mice exhibited the absence of intact parathyroid glands, while retaining normal serum PTH levels, mirroring the characteristics of PT-*Dicer^-/-^* mice. Conversely, PT-*Tsc1^-/-^*mice with hyperactivated mTORC1 exhibited enlarged glands and elevated serum PTH and calcium levels. Significantly, PT-*Dicer^-/-^*;*Tsc1^-/-^* double knockout mice demonstrated a reversal of the aparathyroidism of PT-*Dicer^-/-^* mice, preserving intact parathyroid glands and reinstating CKD-induced SHP. Lastly, data collected from a network of 106 healthcare organizations demonstrated that drug-induced mTOR inhibition is associated with reduced elevation of serum PTH levels in kidney transplant recipients. The latter findings offer physiological validation for our observations in genetically modified mouse models, highlighting the central role of mTORC1 signaling in CKD-SHP. Altogether, our results indicate that mTOR operates downstream of Dicer and miRNA. Consequently, Dicer, miRNA and mTORC1 collectively play a crucial role in maintaining the postnatal integrity and function of the parathyroid glands.

## Introduction

Secondary hyperparathyroidism (SHP) is a common complication of chronic kidney disease (CKD) that correlates with patient morbidity and mortality (1, 2). In experimental SHP, the increase in PTH gene expression is attributed to post-transcriptional mechanisms due to enhanced PTH mRNA stability (3–5). An additional layer of post-transcriptional control is mediated by microRNA (miRNA). miRNA are initially transcribed as longer precursor RNA molecules that are ultimately cleaved by Dicer to form mature and functional miRNA (6). Dysregulated miRNA profiles have been observed in patients with primary hyperparathyroidism, presenting differentially expressed miRNA in parathyroid adenomas and carcinomas when compared to normal parathyroid tissue (7, 8). We have previously demonstrated through miRNA sequencing that human and rodent parathyroids share similar miRNA profiles. Moreover, a significantly disrupted miRNA transcriptome was observed in hyperplastic parathyroid glands from end-stage renal disease patients with SHP, as well as in mice and rats with experimentally-induced CKD-associated SHP (9). Antagonizing two abundant parathyroid miRNA, let-7 and miR-148, led to alterations in serum PTH levels in both controls and CKD rats, along with changes in PTH secretion in isolated parathyroid glands in vitro, pointing to the direct impact of these miRNA on regulation of PTH gene expression (9). To study the global role of miRNA in the parathyroid, we generated mice with targeted gene deletion of *Dicer* and consequently miRNA specifically in their parathyroid glands. Remarkably, PT-*Dicer^-/-^* mice developed normally and had normal basal serum PTH, calcium and phosphate levels. However, PT-*Dicer^-/-^* mice failed to increase serum PTH levels in response to the stimuli of acute and chronic hypocalcemia and CKD, underscoring the central roles of parathyroid *Dicer* and miRNAs in mediating the responses of the parathyroid to the major PTH secretion stimulators (10).

Mechanistic target of rapamycin (mTOR) plays a major role in regulating different cellular processes, including growth and metabolism. mTOR functions as two distinct complexes, mTORC1 and mTORC2 (11). mTORC1 is a key regulator of cell growth and metabolism and exerts its influence by integrating cues such as growth factors, energy levels, cellular stress, and amino acids (12). The promotion of protein synthesis by mTORC1 largely depends on the phosphorylation of two key effectors, ribosomal protein S6 (S6) and eukaryotic translation initiation factor 4E-binding protein (11, 13). Phosphorylation of S6 (pS6) is used as an indicator of mTORC1 activity, playing a beneficial role in protein synthesis and control of cell size. In a prior study, we have demonstrated the activation of parathyroid mTORC1 signaling in SHP. Furthermore, we found that the inhibition of mTORC1 using rapamycin not only prevented but also corrected parathyroid cell proliferation in SHP rats both in vivo and in parathyroid glands in cultures (14). Tuberous sclerosis complex (TSC) genes, *Tsc1 and Tsc2*, encode for hamartin and tuberin proteins, forming the TSC complex that negatively regulates mTORC1. Mutations in either *Tsc1* or *Tsc2* lead to mTORC1 hyperactivation, associated with cellular growth abnormalities and tumorigenesis (15–17).

This study explores the interplay between miRNA and mTORC1 in parathyroid function, employing genetically modified mouse models featuring parathyroid-specific deletions of *Dicer*, *mTORC1* or *Tsc1* (characterized by mTORC1 hyperactivation). Additionally, we studied the combined effects of *Dicer* and *Tsc1* deletions in double knockout mice. Our findings underscore the essential roles of Dicer, miRNA and mTORC1 signaling in maintaining the lifelong integrity and functionality of parathyroid glands, as well as in the context of CKD-induced SHP development.

## Concise methods

### Animal housing

All animal experiments were approved by the Institutional Animal Care and Use Committee of the Hadassah Hebrew University. Animals had free access to food and drinking water.

### Parathyroid-specific knockout mice

Parathyroid-specific knockout mice were generated by Cre-LoxP recombination using PTH-*Cre* mice, which express the *Cre* recombinase under the control of the human parathyroid hormone (PTH) promoter [FVBTg(PTH-Cre); Jackson Laboratory, Bar Harbor, ME] (18), as previously described (10). To visualize the small mouse parathyroid glands using fluorescence microscopy, we introduced EYFP or tdTomato labeling.

Parathyroid-specific *mTORC1* knockout mice (PT-*mTORC1^-/-^*) and Parathyroid-specific *Tsc1* knockout mice (PT-*Tsc1^-/-^*) were generated by crossing PTH-*Cre* mice with mice possessing floxed *mTORC1* (19, 20) *or Tsc1* genes (19, 21), respectively. The appropriate strains were further crossbred to generate the *Dicer* and *Tsc1* double knockout mice and their appropriate controls.

### Genotyping

Total DNA was extracted from ear samples and genotyping was conducted using standard PCR techniques using the appropriate primers (Table 1) (10).

**Table 1.**
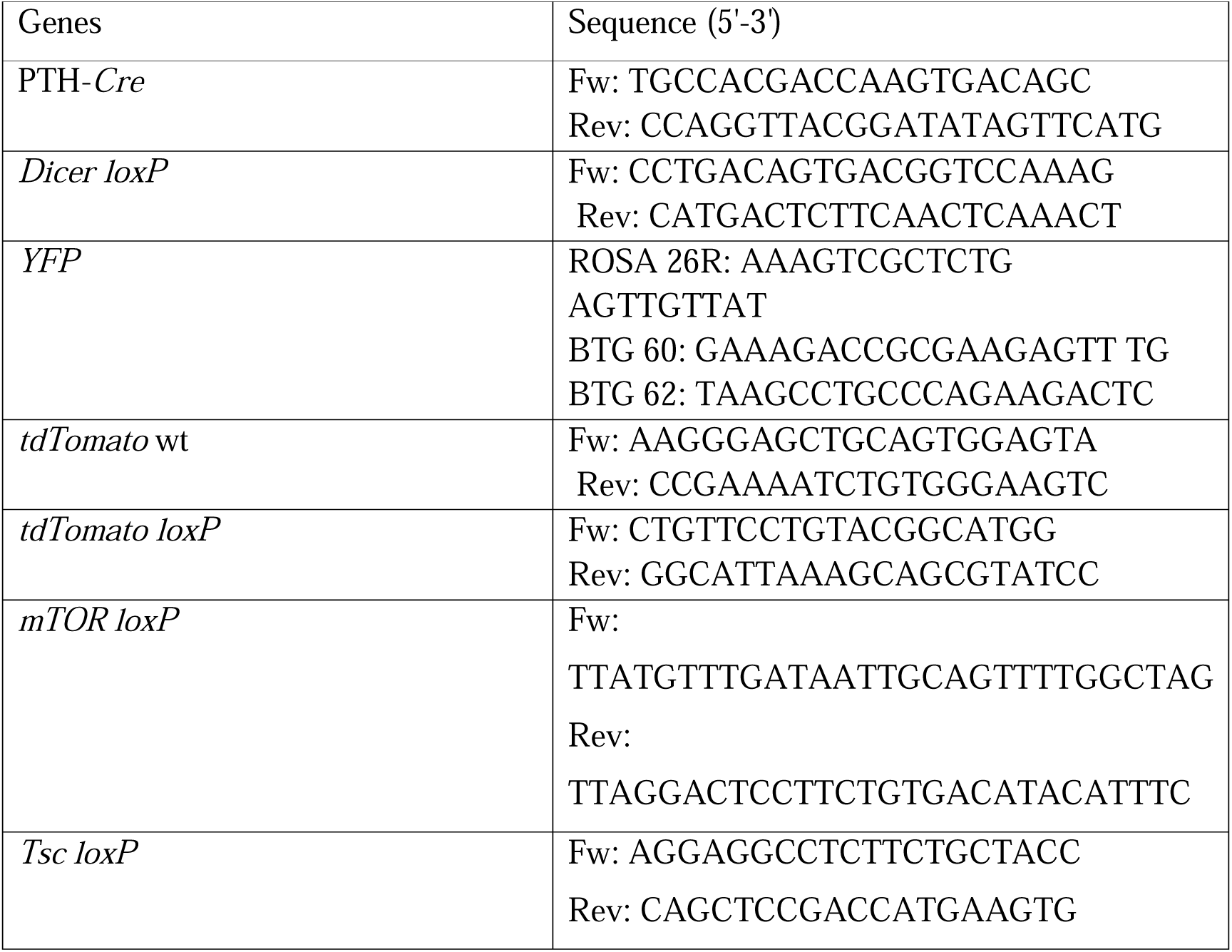
Primers used for DNA genotyping.

### Serum biochemistry and PTH levels

Blood samples were analyzed for serum calcium, BUN and PTH levels (10).

### Thyroparathyroidectomy

Mice were anesthetized and thyroid tissue removed or sham operation performed. At 30 minutes blood samples were collected for PTH.

### RNA isolation and qRT-PCR

RNA from the two parathyroid glands of individual mice was extracted and followed by cDNA synthesis and qRT-PCR with the appropriate priments.

### Microscope fluorescent imaging and quantification

Postnatal mice at various ages were anesthetized, and thyroparathyroids exposed and visualized using GFP or CY3 filters.

### Immunofluorescence staining of mouse paraffin sections

Paraffin mouse thyroparathyroid sections were immunostained using primary antibodies and Fluor chrome–conjugated secondary antibodies Cy3 and Cy5 and visualized by florescence microscope.

### Immunofluorescence staining of mouse embryos

Embryos were obtained from timed matings on specific days of interest and serial sections of paraffin-embedded embryos obtained and stained for PTH (Quidel), CC3 (Cell Signaling Technology), and nuclear staining with DAPI. IHC-generated images, three-dimensional (3D) reconstructions of the parathyroids were created for volumetric data.

### RNAscope in situ hybridization

RNase-free paraffin-embedded sections were studied using specific probes for Gcm2, Pth and Dicer.

### Human clinical investigation

We analyzed data from human kidney transplant recipients undergoing immunosuppressive treatment regimens of mTOR inhibitors or calcineurin inhibitors. Data was collected from the TriNetX network datasets (22) comprising over 127 million individuals, representing data derived from 106 healthcare organizations.

### Statistical analysis

Values are presented as mean±SE. A 2-tailed Student’s t-test was used to assess differences among groups. A p value of less than 0.05 was considered significant.

## Results

### Parathyroid-specific PT-*Dicer^-/-^* mice exhibit undetectable or severely hypoplastic parathyroid glands

It is challenging to accurately identify the minute parathyroid glands embedded within the thyroids, even when utilizing a binocular microscope. To study the parathyroids of PT-*Dicer^-/-^* and PT-*Dicer^+/+^* mice, we therefore initially employed immunohistochemistry (IHC) on serial thyroparathyroid paraffin sections stained for PTH. In most mature control mice, the two parathyroid glands were readily identifiable (**Fig 1A-C**). Missing glands were likely ectopically located, as previously reported in ∼30% of mice (**Fig 1C**) (23). Unexpectedly, intact parathyroid glands could not be discerned in any of the sections from PT-*Dicer^-/-^* mice (n=65 mice) (**Fig 1C**). In the few sections that displayed PTH-producing cells, there were only small, disorganized structures of cell clumps, very different from the intact well-defined parathyroid glands in control mice (**Fig 1A and B**). Histological analysis using H&E staining of parathyroid sections demonstrated the preservation of the parathyroid gland capsule in PT-*Dicer^-/-^* mice, yet adipocytes infiltrated and occupied the void left by the ablated parathyroid cells (**Fig 1B**). It is plausible that in sections from the majority of PT-*Dicer^-/-^* mice, where PTH-producing cells were not detected, the minute cell clumps were too small to be identified by IHC. Therefore, mature PT-*Dicer^-/-^* mice lack intact parathyroid glands and instead exhibit residual parathyroid cell clusters.

**Figure 1.**
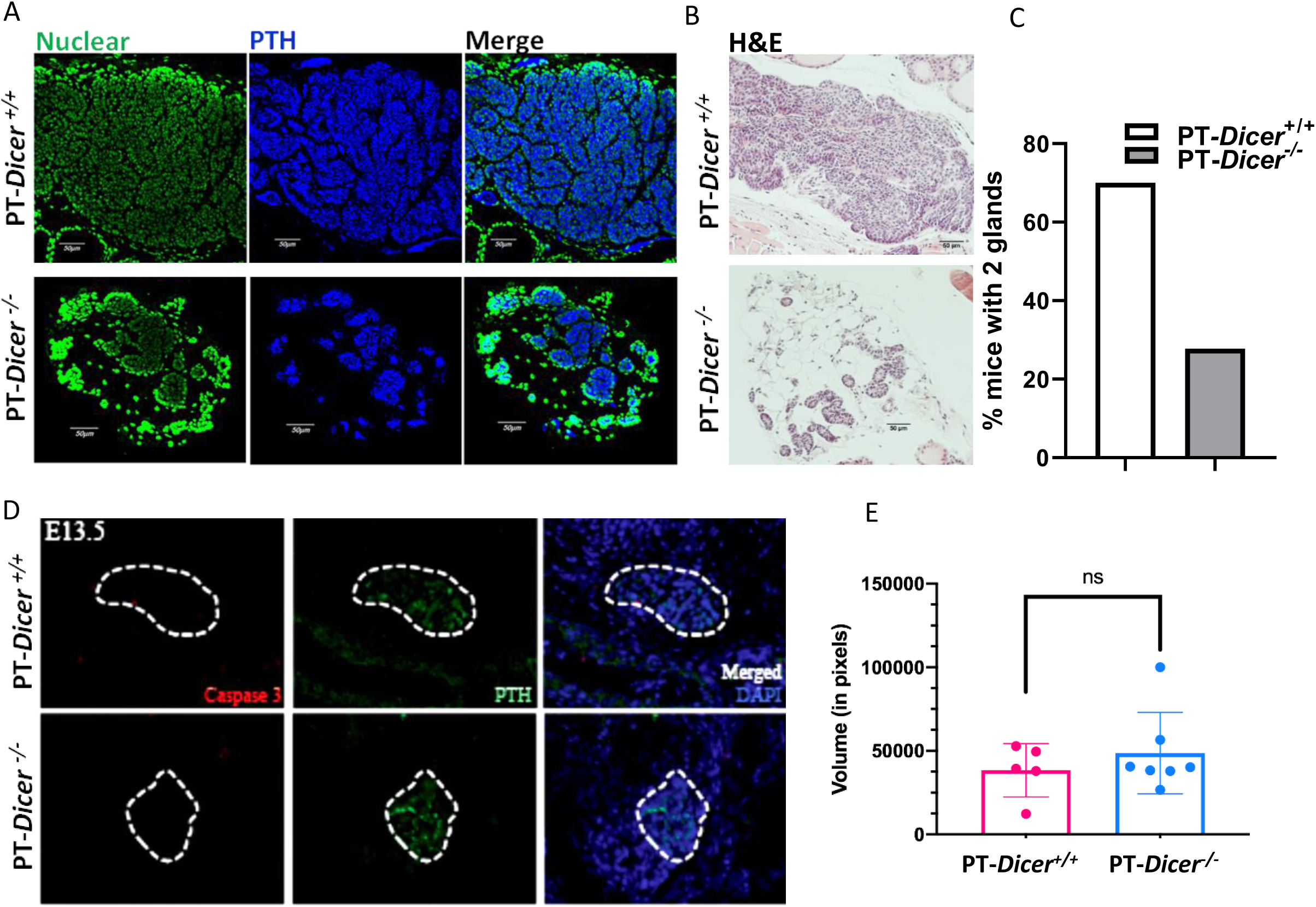
Aberrant parathyroid tissue in mature adult PT-*Dicer*^-/-^ but not in PT-*Dicer^+/+^* embryos. **(A)** Representative immunofluorescence staining of thyroparathyroid sections from control (PT-*Dicer^+/+^*) and parathyroid-specific *Dicer^-/-^*(PT-*Dicer^-/-^*) mice highlighting sections with detectable parathyroid cells, stained for nuclear sytox (green), PTH (blue) and merge images. **(B)** H&E histological sections of thyroparathyroid sctions as above*^/-^* mice. **(C)** Percent of mice with 2 intact parathyroid glands detected in serial sections of thyroparathyroids from PT-*Dicer^+/+^*(n=45) and PT-*Dicer^-/-^* mice (n=65). (**D**) Representative IF staining for PTH and cleaved caspase3 of parathyroids from control PT-*Dicer^+/+^* and PT-*Dicer^-/-^* embryos at E13.5. (**E**) Volumetric analysis of parathyroid embryos as in D (PT-*Dicer^+/+^* n=5, PT-*Dicer^-/-^* n=7 parathyroids).

### Deletion of *Dicer* and consequently miRNA in the parathyroid does not affect the volume of embryonic parathyroid glands or their expression of PTH and caspase3

The initiation of PTH expression starts as early as embryonic day E11.5 (24), implying that expression of the human PTH promoter-driven *Cre* would result in early embryonic *Dicer* disruption in PT-*Dicer*^-/-^ embryos. To follow the development of parathyroid glands during fetal stages, PT-*Dicer^-/-^* and control PT-*Dicer^+/+^*embryos were generated by timed mating and studied at E13.5, E19.5 and E21.5.

Immunofluorescence (IF) staining for PTH in paraffin-embedded whole embryo sections at all embryonic ages consistently revealed two intact parathyroid glands in both PT-*Dicer^-/-^* and control embryos with no difference in PTH staining at all time points. Apoptotic cell death detection by cleaved caspase3 also showed no difference between PT-*Dicer^-/-^* and control mice (**Fig 1D** **and Supplemental Fig 1A&B**).

Likewise, volumetric analysis of the parathyroids demonstrated no disparity (**Fig 1E** and **Supplemental Fig 1C&D**). Hence, our findings indicate that the presence of Dicer and thus miRNA is not imperative for embryonic parathyroid development and for *Pth* embryonic expression, and that the loss of intact parathyroid glands in PT-*Dicer^-/-^* mice occurs after E21.5.

### Parathyroid glands are gradually lost after birth in PT-*YFP;Dicer^-/-^* mice, due to increased apoptosis

To enhance the identification and tracking of parathyroid cells in whole mice, we generated mice with parathyroid-specific YFP labeling using *Cre-Lox* recombination in either control *Dicer^+/+^* or PT-*Dicer^-/-^* mice. PT-*YFP;Dicer^-/-^* mice expressing YFP exhibited normal basal serum levels of calcium, phosphate and PTH, comparable to control PT-*YFP;Dicer^+/+^* mice (**Supplemental Fig 2A-D, control diet**) (10), similar to the findings previously described in PT-*Dicer^-/-^* mice lacking YFP expression (10, 14). Renal failure induced by an adenine-rich high phosphorus diet led to the expected changes in serum biochemistry in both YFP expressing strains (**Supplemental Fig 2A-C**). However, PT-*YFP;Dicer^-/-^* mice did not increase serum PTH levels compared to the expected rise seen in control PT-*YFP;Dicer*^+/+^ mice (**Supplemental Fig 2D**), mirroring the findings in PT-*Dicer^-/-^* mice without YFP expression (10, 14). Therefore, PT-*YFP;Dicer^-/-^* mice manifest the same characteristics as PT-*Dicer^-/-^* mice, which includes normal basal PTH levels and inability to increase serum PTH in CKD.

We then proceeded to monitor the development of parathyroid glands in both control PT-*YFP*;*Dicer^+/+^* and PT-*YFP*;*Dicer^-/-^* mice, following them from birth to adulthood. Fluorescence-guided microscopy of the exposed neck area easily detected two intact parathyroid glands in control mice across all ages, with the majority of glands located near the superior or inferior poles of the thyroid (**Fig 2A&B**). Notably, immediately after birth, at P0 (not shown) and P1, PT-*YFP;Dicer^-/-^* pups also exhibited the presence of two glands (**Fig 2A&B**). However, from P2 onward, a progressive reduction in the intact parathyroid glands was observed in PT-*YFP;Dicer^-/-^* mice, with the left gland persisting longer. By P12, there were no visible intact parathyroid glands in the neck area of PT-*YFP;Dicer^-/-^* mice (**Fig 2A&B**). Therefore, our findings affirm that mice with parathyroid-specific deletion of *Dicer* are initially born with intact parathyroid glands that are subsequently lost shortly after birth, thus confirming the essential role of miRNA postnatally in the preservation of intact parathyroid glands.

**Figure 2.**
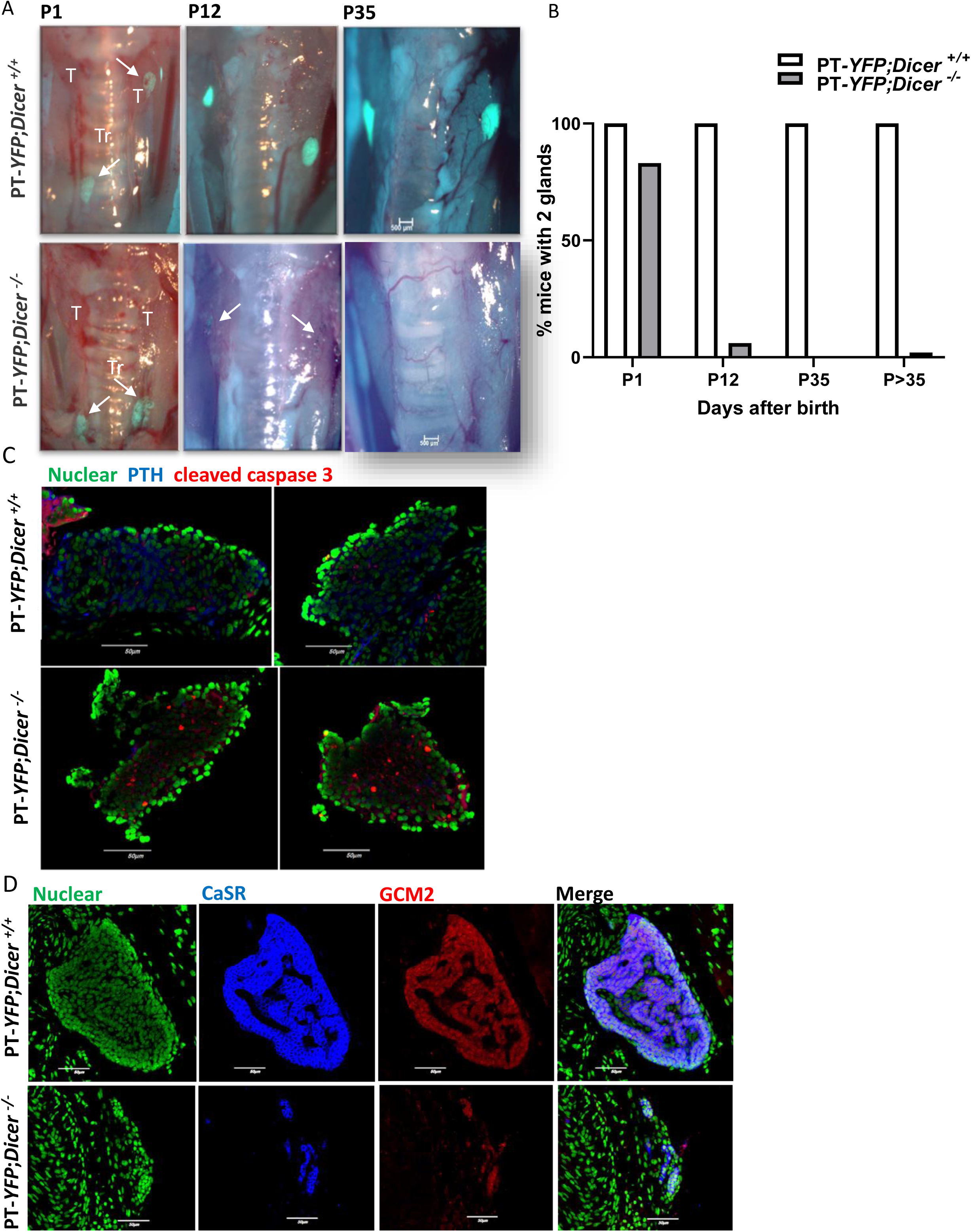
Intact parathyroid glands are gradually lost after birth in PT-*YFP;Dicer^-/-^* mice, as evident by YFP fluorescent microscopy, attributed to increased apoptosis. (**A**) Representative fluorescent microscope images superimposed on bright field images of exposed glands in the neck area of control PT-*YFP;Dicer^+/+^* and PT-*YFP;Dicer^-/-^* mice at postpartum days P1, P12 and P35. The fluorescence intensity exposure was 2 seconds, with a magnification x2. T, thyroid, PT, parathyroid, Tr, trachea. Arrows point to the location of parathyroid glands or cell clusters. (**B**) Percentage of mice with either 2 intact parathyroid glands or no intact glands at each time point as in A (n=6-18) and in adult mice (>35 days to 1 year) (n= 20-44). (**C**) Representative IF staining of paraffin sections from parathyroid glands of P1 (n=2-4) control PT-*YFP*;*Dicer^+/+^* and PT-*YFP*;*Dicer^-/-^* mice stained for cleaved caspase3 (red), PTH (blue) and nuclear sytox (green). (**D**) Representative IF staining of paraffin sections from parathyroid glands of P4 control PT-*YFP*;*Dicer^+/+^* and PT-*YFP*;*Dicer^-/-^* mice stained for CaSR (blue), GCM2(red) and nuclear sytox (green) (n=2-4).

IF staining for cleaved caspase3 of parathyroid gland sections from control PT-*YFP;Dicer^-+/+^* and PT-*YFP;Dicer^-/-^* mice, at P1 and P4, showed increased occurrence of apoptosis in parathyroid glands from PT-*YFP;Dicer^-/-^* neonates. There was no change in staining for PTH or for the specific parathyroid markers glial cells missing 2 (GCM2) and calcium sensing receptor (CaSR) in PT-*YFP;Dicer^-/-^* compared to control PT-*YFP;Dicer^+/+^* pups (**Fig 2C&D**). Taken together, our findings demonstrate that deletion of *Dicer* and the consequent global miRNA deficiency triggers apoptosis in parathyroid cells shortly after birth. This ultimately leads to the loss of intact parathyroid glands in PT-*Dicer^-/-^* mice with no change in serum PTH levels or in the expression of parathyroid markers within the surviving cells.

### The residual parathyroid cell clusters in PT-*Dicer^-/-^* mice still express *Dicer* and adequate PTH production for maintaining basal serum levels

To ascertain the origin of PTH production responsible for maintaining normal serum levels in PT-*YFP;Dicer^-/-^* mice, we performed thyroidectomy in both control and PT-*YFP;Dicer^-/-^* mice. Thyroidectomy removed the thyroid glands along with any associated parathyroid glands in control mice, and the extirpation of any lingering parathyroid cell clusters within PT-*YFP;Dicer^-/-^* mice (**Fig 3A**). As anticipated, thyroparathyroidectomy led to a reduction in serum PTH levels within 30 minutes in control PT-*YFP*;*Dicer^+/+^* mice (25). Intriguingly, a similar decrease in serum PTH levels was observed following thyroidectomy in PT-*YFP;Dicer^-/-^* mice (**Fig 3B**). This result indicates that PTH producing cells are located within the thyroid of these mice and play a substantial role in maintaining basal serum PTH levels in PT-*YFP;Dicer^-/-^*mice, which lack intact parathyroid glands.

**Figure 3.**
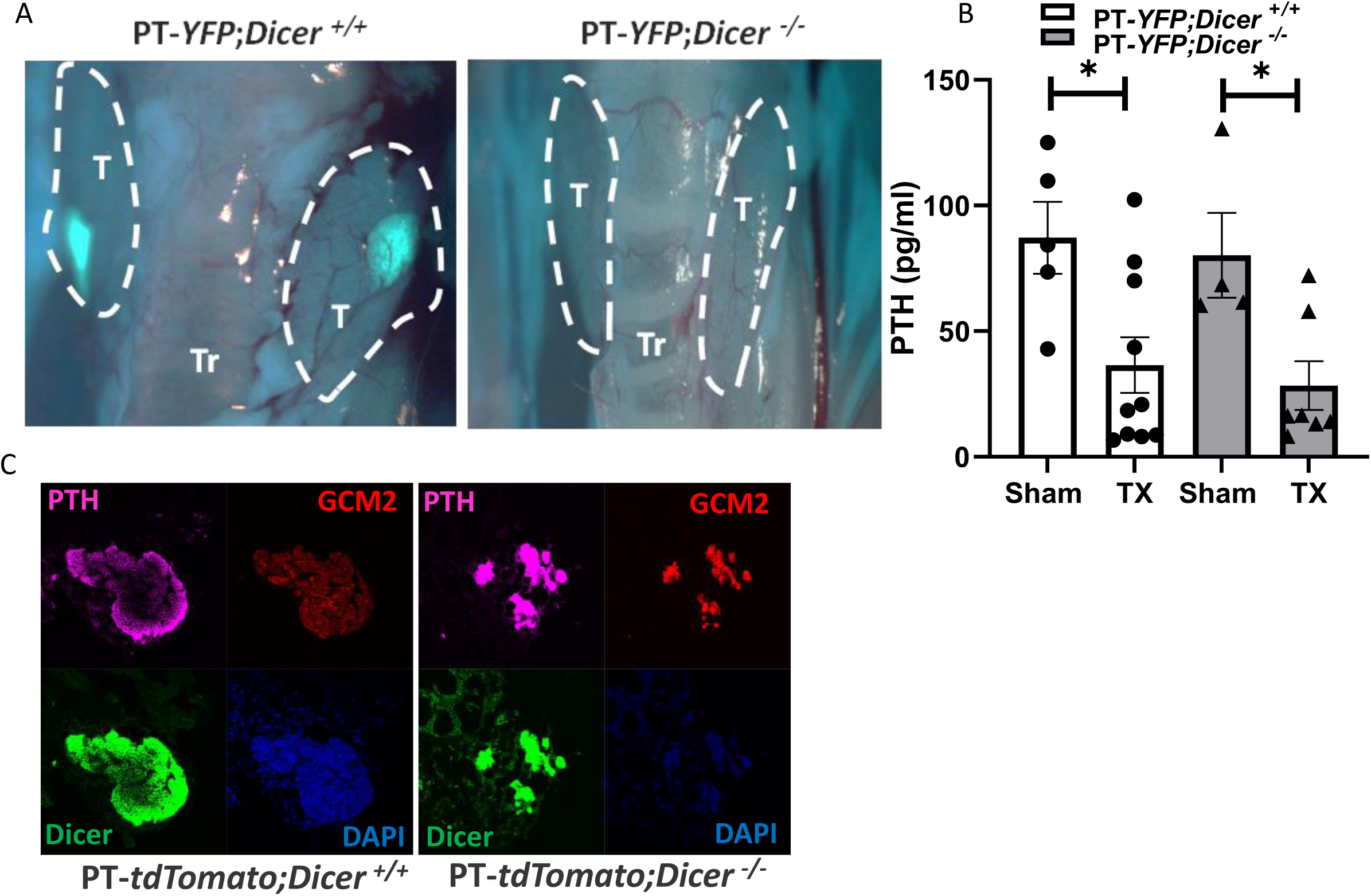
Serum PTH levels are similarly decreased following thyroidectomy in control PT-*YFP;Dicer^+/+^* and PT-*YFP;Dicer^-/-^* mice. (**A**) Thyroidectomy or sham operation was conducted by removal of the thyroid along with embedded parathyroids in control PT-*YFP;Dicer^+/+^* mice, or the thyroids with no intact parathyroid glands in PT-*YFP;Dicer^-/-^* mice, as illustrated by the dashed lines in representative fluorescent dissection microscope images. (**B**) Serum PTH levels 30 min post thyroidectomy or sham operation. Results are presented as mean±SE. *, p<0.05. (**C**) Representative RNAscope analysis of 2-month-old control PT-*tdTomato;Dicer^+/+^* and PT-*tdTomato;Dicer^-/-^* mice depicting the expression of GCM2, Dicer, PTH and nuclear Dapi. n=2.

To characterize the cell clusters still present in PT-*Dicer^-/-^* mice that express and secrete PTH, we performed RNAscope in situ hybridization (ISH). As expected, control PT-*Dicer^+/+^* mice expressed *Pth* and *Gcm2* mRNAs throughout the parathyroid, as well as the targeted *Dicer* sequence. Conversely, most parathyroid cells in PT-*Dicer^-/-^* mice showed no expression for these transcripts at 2 months of age, confirming the successful deletion of *Dicer* and the subsequent absence of intact parathyroid glands. Nevertheless, small cell clusters still retained *Pth*, *Gcm2,* and *Dicer* expression in PT-*Dicer^-/-^* mice (**Fig 3C** and **Supplemental Fig 3**). This suggests that a subset of parathyroid cells evade *Dicer* deletion allowing for their survival while preserving their parathyroid traits and functions. Therefore, the outcome of parathyroid-specific *Dicer* and miRNA ablation is the death of parathyroid cells and loss of parathyroid glands, which is mitigated by a small subpopulation of parathyroid cells evading *Dicer* deletion.

### PT*-tdTomato;Dicer^-/-^* mice reveal the gradual loss of parathyroid glands

The small clusters of parathyroid cells in adult PT-*YFP;Dicer^-/-^* mice were barely identifiable by IF staining and whole mouse parathyroid *YFP* imaging (**Fig 1A&B and** **Fig 2A, P12**). To enhance the visibility of these structures, we generated an additional mouse colony introducing the brighter *tdTomato* fluorescence marker, initially in conjunction with *YFP.* This resulted in parathyroid *Cre-*driven control PT-*YFP;dTomat;Dicer^+/+^* and PT-*YFP;dTomat;Dicer^-/-^* mice. As before, the parathyroids were readily distinguishable in control mice using both *YFP* and *tdTomato* fluorescence guided microscopy, with the *tdTomato* signal being significantly more intense than that of *YFP* (**Fig 4A**). As above, we could not detect intact parathyroid glands in adult PT-*YFP;tdTomato;Dicer^-/-^* mice (**Figs 4A, 2 and 1**). However, with tdTomato imaging, the presence of tiny parathyroid cell clusters, which were not detectable with *YFP*, became more apparent, in PT-*YFP;tdTomato;Dicer^-/-^* mice (**Fig 4A**).

**Figure 4.**
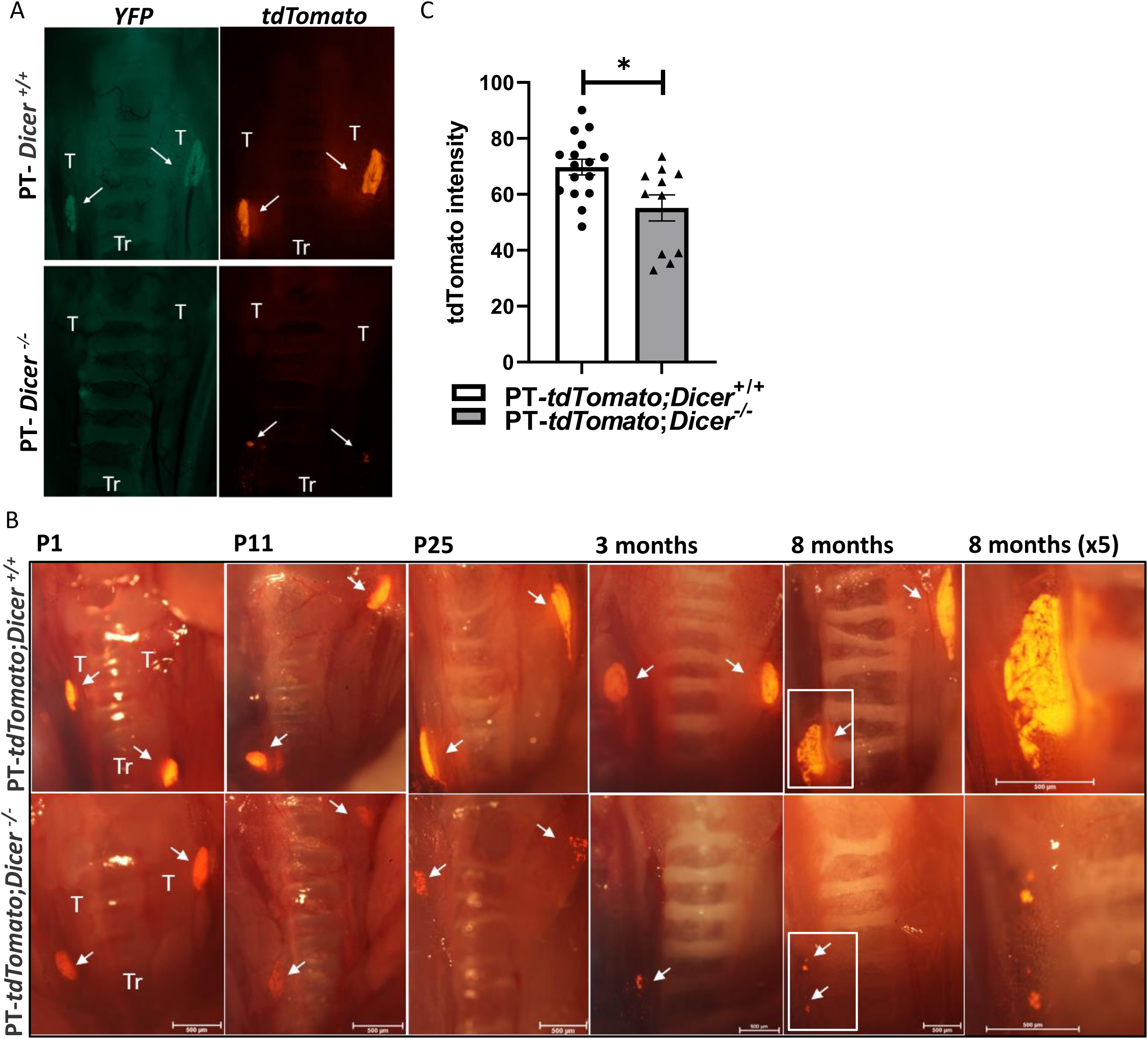
Identification of small parathyroid cell clusters in PT-*tdTomato;Dicer^-/-^*substituting intact glands early after birth. (**A**) Mice engineered to express both *YFP* and *tdTomato* exclusively in the parathyroids were examined by fluorescent guided microscopy of the exposed neck area at 3 months of age. Representative images from dissection microscopy for *YFP* or *tdTomato* display speckles of cell clusters by *tdTomato expression*, but not by *YFP* expression in the PT-*Dicer**^-/-^*** mice. (**B**) Representative fluorescent dissection microscope images superimposed on bright field images of the exposed neck area in postpartum PT-*tdTomato;Dicer^+/+^* and PT-*tdTomato;Dicer^-/-^* mice at various ages. n=2-16. Magnification: P1 x2.5, P11-8 months x2. The right panels additionally exhibit a 5-fold magnification at 8 month. Arrows point to the location of parathyroid glands or cell clusters. T, thyroid; Tr, trachea. Fluorescence intensity exposure was set to 2 seconds for YFP and 0.25 seconds for tdTomato. (**C**) Intensity analysis of the tdTomato signal in parathyroid glands of PT-*tdTomato;Dicer**^+/+^*** control and PT-*tdTomato;Dicer^-/-^* P1 mice as in B, quantified using ImageJ software.

The enhanced intensity of the *tdTomato* signal revealed the timeframe for the progressive degeneration of parathyroid glands in PT-*tdTomato*;*Dicer^-/-^* mice. In PT-*tdTomato*;*Dicer^-/-^* mice, the reduced intensity of the *tdTomato* signal was already visible at P1, when gland size was still comparable to that of control mice (**Fig 4B&C**). Remnants of parathyroid glands were still visible even at P25 with the tdTomato reporter, although exhibiting diminished density and a more punctate morphology when contrasted with the parathyroids of control mice (**Fig 4B**). Taken together, these findings show that PT-*Dicer^-/-^* mice are born with intact parathyroid glands that are gradually lost after birth. Parathyroid cell clusters that still express *Dicer* maintain normal serum PTH levels in PT-*Dicer^-/-^* mice.

### Reduced parathyroid mTORC1 pathway activity in PT-*Dicer^-/-^* mice

We have previously demonstrated the activation of the parathyroid mTORC1 pathway in rats with CKD-SHP. Additionally, mTORC1 inhibition by rapamycin prevented and corrected parathyroid cell proliferation in these rats (14). We therefore assessed the impact of *Dicer* and miRNA ablation on the parathyroid mTORC1 pathway. Indeed, pS6 (Ser235/236), a marker for mTORC1 activity, decreased in parathyroid, the stage when the parathyroid glands were still intact (**Fig 5A-B**). A similar decrease in pS6 was observed by IF staining of parathyroid sections of adult PT-*Dicer^-/-^* compared to control mice, showing the reduction in pS6 staining within the remaining parathyroid cell clusters (**Fig 5C-D**). Hence, deletion of *Dicer* and miRNA results in reduced parathyroid mTORC1 pathway activity, suggesting that *Dicer* and miRNA ablation could affect the mTORC1 pathway in PT-*Dicer^-/-^* mice.

**Figure 5.**
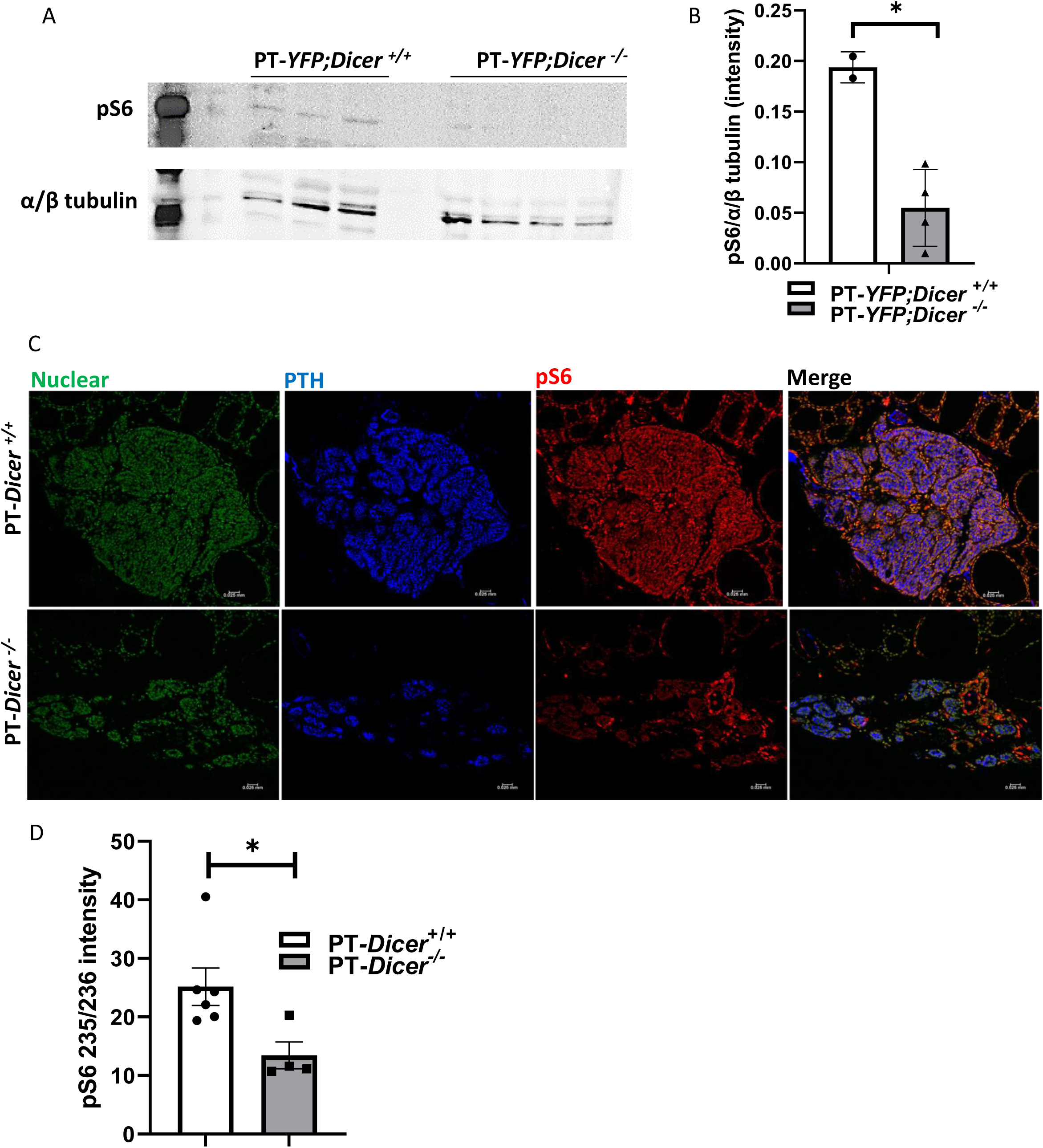
Parathyroid-specific PT-*Dicer^-/-^* mice have reduced phosphorylated S6 in their parathyroids. (**A**) Representative Western blot showing pS6 (Ser235/236) (pS6) and α/β tubulin proteins from P1 PT-*YFP*;*Dicer^+/+^* and PT-*YFP*;*Dicer^-/-^* mice. Protein extracts are from pooled microdissected parathyroid glands of 8-12 mice in each lane. (**B**) Quantitative assessment of pS6 normalized to α/β tubulin as shown in A. (**C**) Representative IF staining of paraffin sections from parathyroid glands of 3 month-old control PT-*Dicer^+/+^* and PT-*Dicer^-/-^* mice stained for nuclear sytox (green), PTH (blue) and pS6 (red). (**D**) Quantification of pS6 intensity, as shown in C, quantified using ImageJ software. *, p<0.05.

### PT-*mTORC1^-/-^* mice lack intact parathyroid glands from early after birth, yet maintain normal serum PTH levels, similar to PT-*Dicer^-/-^* mice

We generated mice with parathyroid specific ablation of m*TORC1* and *td-Tomato* expression (PT-*mTORC1^-/-^* mice). The PT-*mTORC1^-/-^* mice developed normally, and were viable and fertile. Survival of the PT-*mTORC1^-/-^* mice was similar to that of control littermates, as measured up to 1 year (not shown). There were also no differences in serum calcium (0.60 ±0.01 mmol/L in control and 0.58 ±0.03 mmol/L in PT-*mTORC1^-/-^* mice, n=12) and phosphate levels (0.85 ±0.07 mmol/L control and 0.74 ±0.04 mmol/L PT-*mTORC1^-/-^* mice, n=5) in PT-*mTORC1^-/-^* mice compared to control PT-*mTORC1^+/+^* mice, both at 3 months and other time points (data not presented). Serum PTH levels were similar to controls at all ages tested (**Fig 6C**).

**Figure 6.**
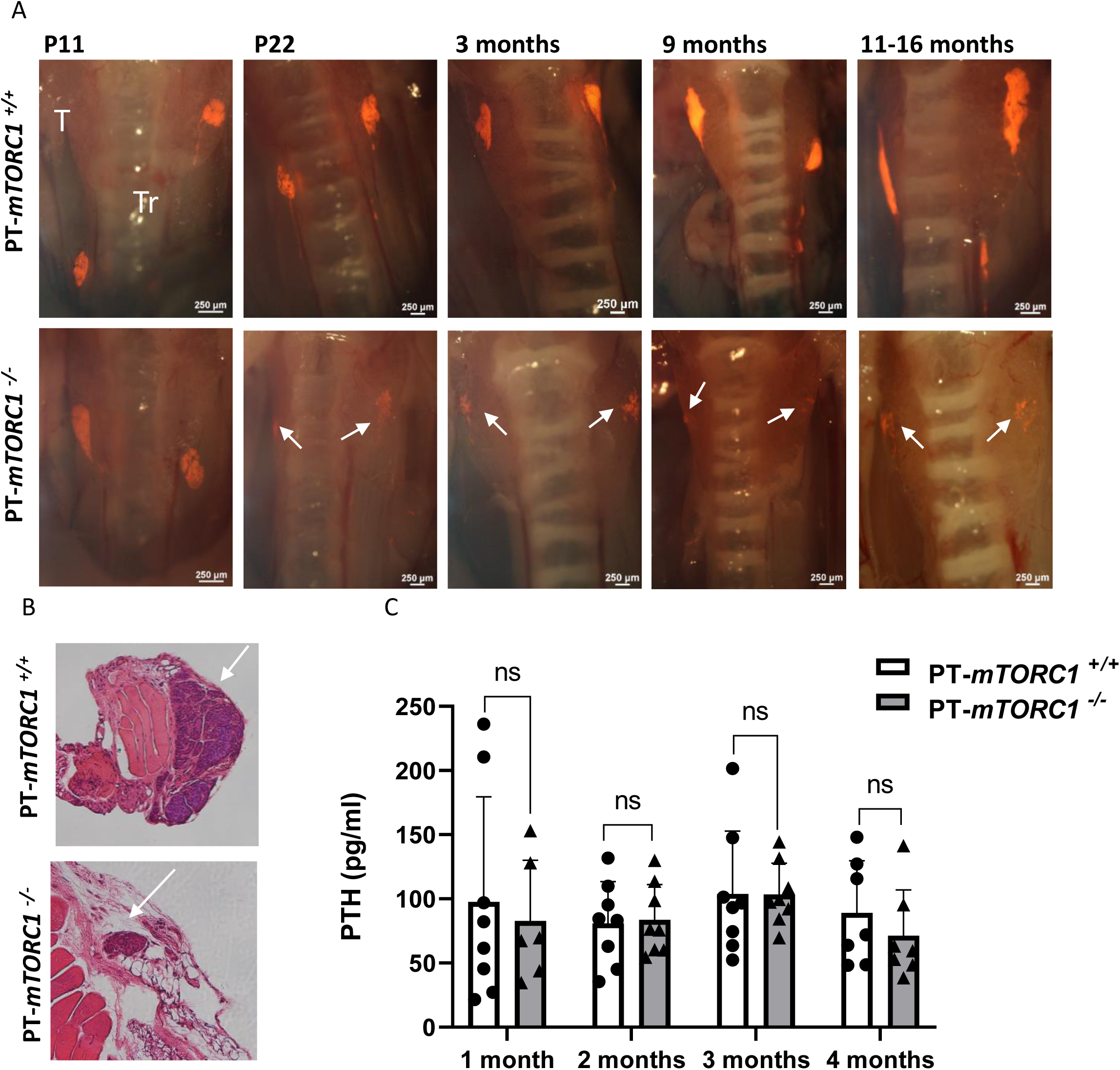
Parathyroid-specific *mTORC1^-/-^* mice exhibit disrupted parathyroid glands from early after birth yet maintain normal serum PTH levels. (**A**) Representative fluorescent microscopy images of tdTomato fluorescence, superimposed on bright field images of the exposed parathyroids from control PT-*mTORC1^+/+^*and PT-*mTORC1^-/-^* mice at various postnatal stages. n=6-16. T, thyroid; Tr, trachea. Arrows point to the parathyroid glands or residual cell clusters. Fluorescence intensity exposure was set to 67 milliseconds and the magnification x2. (**B**) H&E histological sections from PT-*mTORC1^+/+^* and PT-*mTORC1^-/-^* mice. (**C**) Serum PTH levels in PT-*mTORC1^+/+^* and PT-*mTORC1^-/-^* mice at various ages.

Fluorescence guided microscopy showed two intact parathyroid glands in PT-*mTORC1^-/-^* mice at P11 and at earlier stages (**Fig 6A** and not shown), similar to control PT-*mTORC1^+/+^* mice. However, from P11 onward, a progressive reduction in gland size ensued, characterized by a punctuated appearance. Ultimately, fully adult PT-*mTORC1^-/-^* mice had no intact glands, despite normal serum biochemical profiles and PTH levels (**Fig 6A&C**). The remnant cell clusters in PT-*mTORC1^-/-^* mice were also evident by H&E staining of thyroparathyroid sections (**Fig 6B**). There was no change in PTH content in the remnant parathyroids of PT-*mTORC1^-/-^* compared to the intact glands of control mice as indicated by the intensity of IF staining in thyroparathyroid sections (**Supplemental Fig 4A&B**), presumably because PTH was actively secreted to compensate for the decreased parathyroid cell mass. Interestingly, there was a decrease in PTH mRNA with no change in CaSR mRNA in parathyroids of PT-*mTORC1^-/-^* mice at 3 and 4 months (**Supplemental Fig 5** and not shown). Hence, *mTORC1* ablation yields smaller, disrupted parathyroid tissue soon after birth. Remarkably, despite the lack of intact glands and reduction in the number of PTH producing cells, PT-*mTORC1^-/-^* mice maintain normal serum PTH levels, mirroring the phenotype seen in PT-*Dicer^-/-^* mice in the context of *Dicer* ablation.

### Parathyroid-specific *Tsc1^-/-^* mice exhibit enlarged parathyroid glands and increased serum PTH and calcium levels

To gain deeper insights into the contribution of *mTORC1* in the parathyroid, we generated mice with parathyroid-specific mTORC1 hyperactivation through ablation of the mTOR repressor, TSC1, combined with tdTomato expression. mTORC1 was indeed activated in the parathyroids of PT-*Tsc1^-/-^* mice, as indicated by increased pS6 IF staining (**Supplemental Fig 6**). PT-*Tsc1^-/-^* mice developed normally and did not display gross physical abnormalities. However, there was a pronounced reduction in the survival rate of PT-*Tsc1^-/-^* mice when compared to their control littermates, as observed from 1 month to 1 year (**Fig 7A**). Unlike controls, only a single PT-*Tsc1^-/-^* mouse survived up to 11 months. Serum PTH levels in PT-*Tsc1^-/-^* mice were increased 1.5-fold at 1 month when the glands were already enlarged, and continued to increase with age (**Fig 7B**). Serum calcium levels were increased at both 3 and 5 months consistent with the high serum PTH levels (**Fig 7C**). The premature mortality observed in PT-*Tsc1^-/-^* mice is likely attributed to the hyperparathyroidism-induced hypercalcemia, similar to the increased mortality in primary hyperparathyroidism patients with increased serum calcium levels (26).

**Figure 7.**
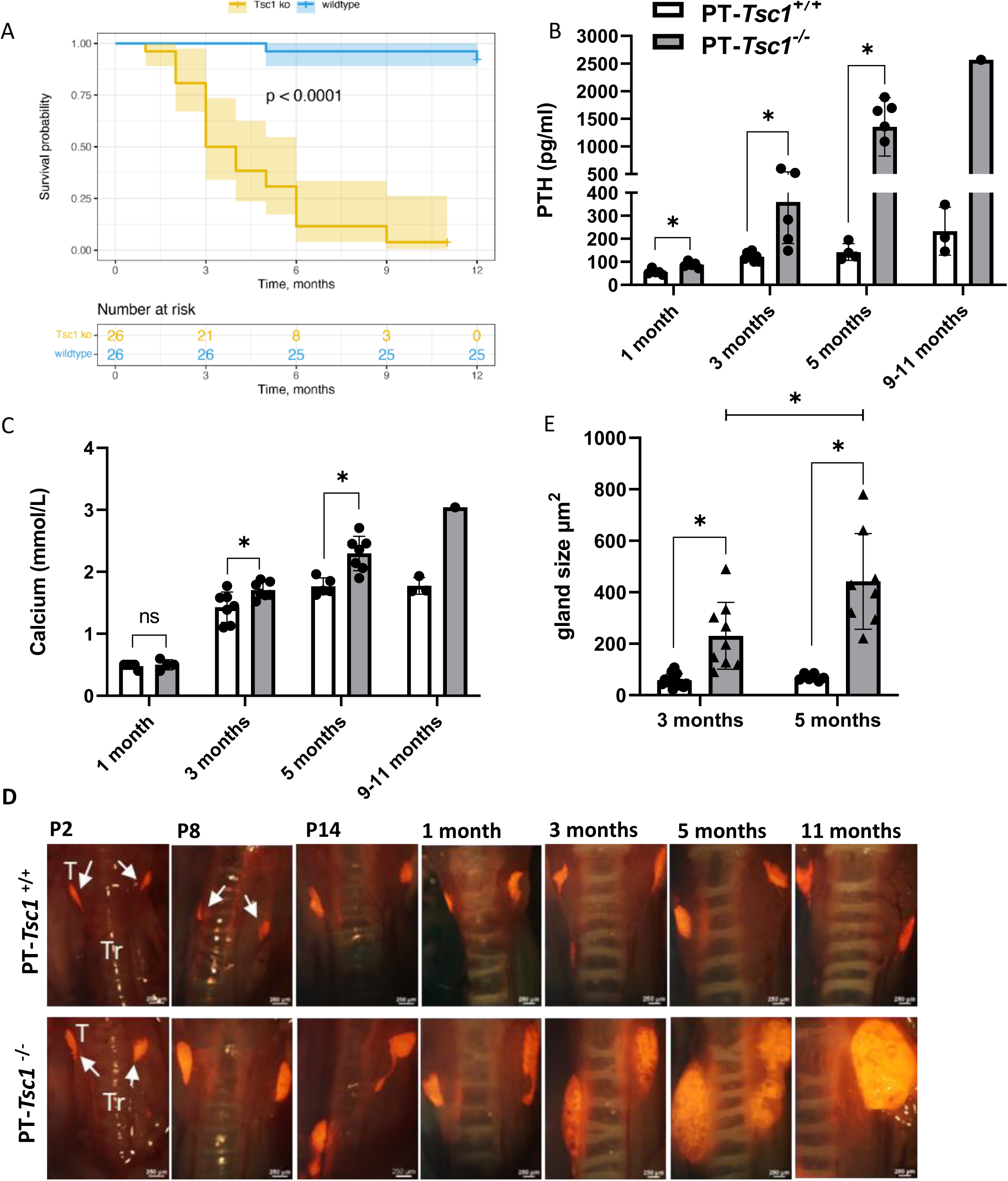
PT-*Tsc1^-/-^* mice have decreased survival, increased serum PTH and calcium levels and enlarged parathyroid glands from early after birth. (**A**) Survival probability curves based on Kaplan-Meier estimates of control PT-*Tsc1^+/+^* ((blue) and PT-*Tsc1^-/-^* (orange) mice at 1 month to 1 year. (**B**) Serum PTH at different ages. (**C**) Serum calcium measured using a Atellica™ CH Calcium assay. Only a single PT-*Tsc1^-/-^* mouse that survived to 11 months is shown. (**D**) Representative fluorescence microscopy images for tdTomato, laid over bright field images of the exposed parathyroids from control PT-*Tsc1^+/+^* and PT-*Tsc1^-/-^* mice at different postnatal ages. n=3-14. T, thyroid; Tr, trachea. The fluorescence intensity exposure was 67 milliseconds and the magnification x1.9. The image of the PT-*Tsc1^-/-^* mouse at 11 month shows only one of the 2 parathyroids at this magnification, due to their large size. (**E**) Parathyroid gland size as measured using ImageJ platform. *, p<0.05.

Fluorescence guided microscopy in PT-*Tsc1^-/-^* mice revealed parathyroid glands of similar size to those of control PT-*Tsc1^+/+^* mice at P2. However, from P8 onward, PT-*Tsc1^-/-^* pups had visibly larger parathyroid glands compared to controls, which persisted and escalated substantially over time (**Fig 7D** **and Supplemental Fig 7**). Parathyroid gland 2-dimensional measurements reaffirmed these results. At 3 months, parathyroid glands in PT-*Tsc1^-/-^* mice displayed a 4-fold increase in size compared to those of control PT-*Tsc1^+/+^* mice. This enlargement continued to escalate, reaching 6 times the size of controls at the 5-month (**Fig 7E** **and Supplemental Fig 8**).

PTH mRNA levels were increased in PT-*Tsc1^-/-^* mice with an in-consistent increase in CasR mRNA (**Supplemental Fig 9**). Of interest, the rise in PTH mRNA corresponded to the elevation in serum PTH levels, although it remained notably milder than the evident enlargement in gland size. Serum PTH levels also only partially mirrored the substantial growth in gland size (**Fig 7B&E**). Taken together, hyperactivation of mTORC1 in PT-*Tsc1^-/-^* mice leads to a significant enlargement of the parathyroid glands postnataly, coupled with increased serum PTH and calcium levels, underscoring the pivotal role of mTORC1 in parathyroid gland morphology and parathyroid cell survival.

### Parathyroid-specific *Dicer* and *Tsc1* double knockout reverses the aparathyroidism of PT*-Dicer^-/-^* mice and restores their CKD induced SHP

To further affirm the miRNA-mTORC1 axis in the parathyroid, we generated parathyroid specific double knockout mice with ablation of both *Dicer* and *Tsc1* coupled with *td-Tomato* expression (PT-*Dicer^-/-^;Tsc1^-/-^* mice). Remarkably, PT-*Dicer^-/-^;Tsc1^-/-^* mice exhibited intact parathyroid glands comparable in size to those of control mice, in contrast to PT-*Dicer^-/-^* mice. This observation was consistent at 2 months of age and across all ages examined up to 9 months (**Fig 8A** and not shown). Serum PTH levels in PT-*Dicer^-/-^*;*Tsc1^-/-^* mice were similar to those of control mice. Importantly, the simultaneous ablation of both *Dicer* and *Tsc1* not only salvaged the structural integrity of the parathyroid glands, but also reinstated their capacity to elevate serum PTH levels in response to adenine high-phosphorus-induced CKD (**Fig 8B**). Hence, the findings from PT-*Dicer^-/-^*;*Tsc1^-/-^* mice underscore that *Tsc1* ablation effectively overrides the lack of parathyroid glands observed in PT-*Dicer^-/-^* mice. This highlights the hierarchical relationship between Dicer and miRNA ablation and their subsequent influence on mTORC1 activity downstream, affecting parathyroid gland integrity and function.

**Figure 8.**
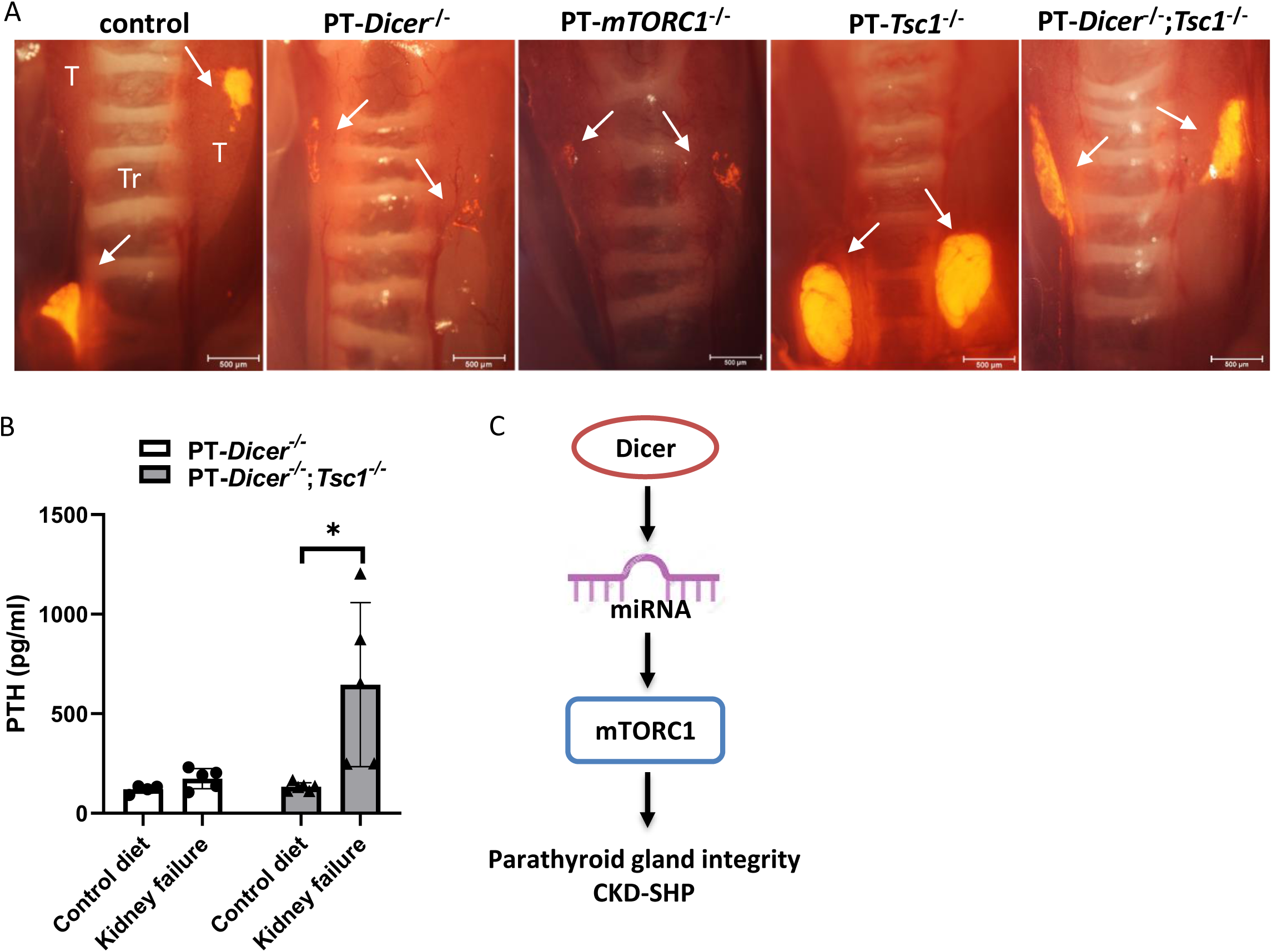
Parathyroid-specific *Dicer* and *Tsc1* double knockout (PT-*Dicer^-/-^*;*Tsc1^-/-^*) reverses the aparathyroidism of PT*-Dicer^-/-^* mice and restores their ability to increase PTH after CKD. (**A**) Representative fluorescent microscopy images of tdTomato fluorescence superimposed on bright field images of parathyroid glands of 2 month-old control (PT-*Dicer^+/+^*;*Tsc1^+/+^*) mice and mice with and parathyroid-specific ablation of *Dicer, mTORC1, Tsc1*, *or Tsc1* and *Dicer* double ablation. Exposure was 250 milliseconds. Arrows indicate parathyroid glands or cell clusters, T, thyroid, Tr, trachea. (**B**) Serum PTH levels in either PT-*YFP*;*Dicer^-/-^* or PT-*Dicer^-/-^*;*Tsc1^-/-^* mice that were fed a control or an adenine-rich high phosphorus diet for 3 weeks to induce CKD and SHP. Results are presented as mean±SE. *, p<0.05 compared to mice fed a control diet. (**C**) Model outlining the relationship between Dicer, miRNA and the mTOR pathway. In normal basal conditions, Dicer facilitates miRNA maturation contributing to basal mTORC1 pathway activity, which is essential for maintaining intact parathyroid glands and the development of CKD induced SHP.

### Drug-induced mTOR inhibition and reduced serum PTH levels in kidney transplant CKD recipients

To examine the physiological relevance of our findings, we studied serum PTH levels in kidney transplant patients whose immunosuppressive regimen is based on mTOR inhibitors compared to patients treated with calcineurin inhibitor-centered regimen. Our study involved data collected from a comprehensive pool of 106 healthcare organizations through the TriNetX network. We established two distinct cohorts for comparative analysis, both comprising patients who had received kidney transplants within a period of 1 to 2 years. The first cohort included individuals with a medication code for “calcineurin inhibitors” (CNI), with the exclusion of those who had a concurrent medication code for mTOR inhibitors. The second cohort consisted of patients with a medication code for “mTOR inhibitors” (mTORi), excluding those with a concurrent medication code for calcineurin inhibitors. Notably, we observed that the mean serum PTH level in patients treated with mTOR inhibitors was significantly lower than that in patients treated with calcineurin inhibitors, registering at 103 pg/ml versus 161 pg/ml (p<0.0001). This discrepancy held true despite both groups displaying similar mean levels of creatinine, urea nitrogen, and phosphorus, with mTOR inhibitor-treated patients exhibiting slightly lower total calcium levels.

Additionally, a smaller percentage of patients in the mTOR inhibitor cohort were diagnosed with hyperparathyroidism, standing at 13.5% as opposed to 24.7% in the calcineurin inhibitor-treated group (p<0.0001), as outlined in **Table 2**. Thus, mTOR inhibition by immunomodulatory medications possibly reduces serum PTH levels in transplant patients. In experimental CKD models, rapamycin corrected and prevented parathyroid cell proliferation both in vivo and in parathyroid glands in cultures (14), whereas reduced calcineurin activity is associated with increased PTH (27). Collectively, our findings underscore the pivotal roles of parathyroid mTORC1 signaling in the development of CKD-associated SHP.

**Table 2:** Demographic, clinical and laboratory characteristics of the two defined kidney transplant patient cohorts, as compared and downloaded from TriNetX on 5/Sep/2023.

| Characteristic | Code | Cohort 1 (CNI)<br>N=42,782 | Cohort 2<br>(mTORi)<br>N=430 | P-value |
| --- | --- | --- | --- | --- |
| Age, years |  | 55.8±15.5 | 57.7±17.5 | 0.0096 |
| Sex, male |  | 58.0% | 62.1% | 0.0862 |
| Hyperparathyroidism | ICD-10 E21.3 | 24.7% | 13.5% | <0.0001 |
| Parathyroidectomy | CPT 1014226 | 2.3% | 2.3% | 0.9486 |
| Cinacalcet | RxNorm 407990 | 22.4% | 8.1% | <0.0001 |
| PTH, pg/ml | TNX 9039 | 161±202<br>(N=31,653) | 103±108<br>(N=251) | <0.0001 |
| Creatinine, mg/dl | TNX 9024 | 1.85±2.02<br>(N=39,884) | 1.90±2.06<br>(N=374) | 0.6714 |
| Phosphate, md/dl | TNX 9027 | 3.37±1.01<br>(N=37,910) | 3.34±0.91<br>(N=331) | 0.5540 |
| Calcium, mg/dl | TNX 9022 | 9.41±0.70<br>(N=39,820) | 9.26±0.71<br>(N=375) | <0.0001 |
| Magnesium. mg/dl | TNX 9026 | 1.78±0.31<br>(N=35,516) | 1.96±0.28<br>(N=281) | <0.0001 |
| Urea nitrogen, mg/dl | TNX 9030 | 27.2±16.3<br>(N=39,254) | 27.0±18.9<br>(N=367) | 0.8007 |
| Bicarbonate, mmol/l | TNX 9021 | 24.2±3.6<br>(N=40,194) | 24.2±3.8<br>(N=383) | 0.8656 |
| Albumin, g/dl | TNX 9045 | 3.92±0.59<br>(N=38,242) | 3.72±0.65<br>(N=350) | <0.0001 |
| Hematocrit, % | TNX 9013 | 37.5±8.8<br>(N=40,338) | 36.5±9.2<br>(N=384) | 0.0411 |

## Discussion

The present study underscores the pivotal roles of Dicer, miRNA, and the mTORC1 signaling pathway in maintaining intact parathyroid glands throughout mouse lifespan. Our prior work utilizing parathyroid-specific *Dicer* knockout mice (PT-*Dicer^-/-^*), illuminated the importance of Dicer and miRNA in stimulating the parathyroid during both acute and chronic hypocalcemia and uremic conditions (10). Building on this PT-*Dicer^-/-^* mouse model, we now reveal an unexpected consequence of *Dicer* and miRNA deficiency-the postnatal apoptosis-driven loss of intact parathyroid glands shortly after birth, leaving only scattered clumps of PTH-producing cells in adult mice. This intriguing revelation underscores the non-essential roles of Dicer and miRNA in embryonic parathyroid development, while emphasizing their vital function in preserving postnatal and lifelong structural integrity of the parathyroid glands. As previously reported, serum PTH levels in these mice remained within the normal range, suggesting that the dispersed cell clusters adequately produce PTH to maintain basal PTH levels.

The transcription factor *Gcm2*, is expressed during both developmental stages and in the adult (28). In humans, *GCM2* loss of function mutations cause idiopathic and familial hypoparathyroidism (29). Various mouse models have shown that *Gcm2* null mutations cause aparathyroidism in the fetal and adult mouse (30–32). As opposed to *GCM2*-knockout mice in which parathyroid cells undergo apoptosis by E12 (31), PT-*Dicer^-/-^* mice described in this study are born with intact glands that undergo apoptosis shortly after birth, gradually leading to eventual absence of parathyroid glands post day P12. We employed 3 mouse strains of parathyroid-specific *Dicer* ablation: PT-*Dicer^-/-^*, PT-*YFP;Dicer^-/-^*, and PT-*tdTomato;Dicer^-/-^*, along with respective control mice to demonstrate that adult PT-*Dicer^-/-^* mice across all strains lack intact parathyroid gland that are replaced by scattered clumps of parathyroid cells in the thyroid tissue or neck area that are more effectively detected using the sensitive tdTomato marker. Notably, *GCM2* expression remained unchanged in the parathyroids of PT-*Dicer^-/-^* mice. These parathyroid cell clusters appear sufficient to maintain normal basal PTH and calcium levels in PT-*Dicer^-/-^*mice, even in the absence of intact parathyroid glands. Thyroidectomy in PT-*Dicer^-/-^* mice led to decreased PTH levels similar to control mice, where intact parathyroid glands were removed along with the thyroid. The origin of the cell clusters in PT-*Dicer^-/-^* mice remains unclear. RNAscope analysis indicated that these cells still express *Dicer*, *Pth*, and *Gcm2*. The incomplete *Cre*-mediated *Dicer* deletion may arise from variable *Cre* expression or its expression in distinct parathyroid cell populations at different time points. The gradual loss of cells postnatally could be because of different cell populations having different requirements for Dicer, or reflect gradual turnover of cells as they progress through a sensitive part of the cell cycle.

Consequently, only cells failing to delete *Dicer* survive, retaining parathyroid identity and function. The outcome of *Dicer* and miRNA ablation therefore is death of parathyroid cells, resulting in a condition of aparathyroidism. The survival of some cells post-apoptosis could explain the milder phenotype compared to complete parathyroid cell loss due to *Dicer* ablation. These scattered clusters seemingly provide sufficient basal serum PTH through excessive PTH production, potentially rendering them unresponsive to further stimulation by hypocalcemia or CKD (10). Alternatively, the impaired structure or smaller cell mass of the hypoplastic parathyroids may hinder their ability to further elevate serum PTH levels. Indeed, studies have demonstrated the critical role of intact parathyroid tissue architecture in the direct effect of high phosphate on stimulating PTH secretion (33, 34).

Aberrant miRNA regulation has been associated with various parathyroid-related diseases, such as primary hyperparathyroidism and CKD-induced SHP. Altered miRNA profiles have been detected in parathyroid adenomas and carcinomas, implying their potential role in these conditions (8). Through parathyroid miRNA deep sequencing, we have previously shown shared profiles between human and rodent parathyroids. Moreover, there were altered miRNA profiles in parathyroid glands from uremic and normal rats as well as in glands form CKD-related SHP patients compared to patients with normal renal function. The manipulating of specific dysregulated miRNAs in experimental SHP using antagomir oligonucleotides has been shown to modify PTH secretion both in vivo and in vitro (9). Evolutionary conservation of miRNAs in normal parathyroid glands and their dysregulation in SHP underscore their significance in parathyroid function and hyperparathyroidism development (9, 35). Here we identify the mTOR pathway as crucial for parathyroid gland morphology and function.

The interaction between miRNAs and the mTOR pathway, wherein miRNAs can either inhibit or stimulate mTOR signaling, is well documented (36, 37). For example, in Xenopus development, inhibiting miRNA biogenesis resulted in reduced size of the pronephric kidney, attributed to decreased mTORC1 activity (36, 38). Notably, the involvement of mTORC1 signaling extends to the realm of parathyroid gland biology. Genomic alterations in the PI3K/AKT/mTOR and Wnt signaling pathways have been associated with parathyroid carcinoma (39–41). Moreover, mutations in the *Tsc1* gene, leading to mTORC1 hyperactivation, correlate with enlarged parathyroid glands and elevated serum PTH levels (42, 43). We have now exposed a significant association between mTOR inhibition in kidney transplant recipients and lower PTH levels. In experimental rat models of CKD-induced SHP, parathyroid mTORC1 signaling activation has been documented. Pharmacological inhibition of mTORC1 effectively prevented and reversed parathyroid cell proliferation, while also halting the increase in PTH secretion in these rats (14).

We have now demonstrated that loss of *Dicer* in PT-*Dicer^-/-^* mice leads to a decrease in mTORC1 activity, suggesting a sequential relationship between the two. Indeed, PT-*mTORC1^-/-^* mice lacked intact parathyroid glands after birth, but preserved normal PTH levels, comparable to the phenotype of *PT-Dicer^-/-^* mice. Conversely, PT-*Tsc1^-/-^* mice with mTORC1 hyperactivation exhibited enlarged glands and increased serum PTH and calcium levels. These findings underscore the pivotal role of the mTORC1 pathway in upholding the structural integrity of the parathyroid glands. The precise mechanism through which mTOR signaling influences parathyroid gland integrity and the survival of parathyroid cells remains to be elucidated. Importantly, in PT-*Dicer^-/-^;Tsc1^-/-^* mice with dual *Dicer* and *Tsc1* ablation, intact glands were preserved, overriding the aparathyroidism state associated with *Dicer* deletion. Moreover, unlike PT-*Dicer^-/-^* mice, PT-*Dicer^-/-^;Tsc1^-/-^* mice demonstrated increased serum PTH levels in the context of CKD-induced SHP. Thus, the PT-*Dicer^-/-^*;*Tsc1^-/-^* mice demonstrate that *Tsc1* ablation effectively supersedes the absence of parathyroid glands seen in PT-*Dicer^-/-^* mice. This observation further underscores the hierarchical relationship between Dicer, miRNA, and mTORC1, implying that Dicer and miRNA operate upstream of mTORC1. Collectively, our results indicate that the ablation of *Dicer* and miRNA leads to a reduction in mTORC1 pathway activity, ultimately culminating in the demise of parathyroid glands (**Fig 8C**). Dicer, miRNA, and mTORC1 emerge as vital contributors to the maintenance of parathyroid gland integrity postnatally, as well to the development of CKD-induced SHP, shedding light on their importance in parathyroid cell survival and function and highlighting new avenues for understanding and potentially treating parathyroid-related disorders.

## Supporting information

Supplemental Mathods

## Acknowledgments

We thank Oded Volovelsky and Moris Nechama (Department of Pediatric Nephrology, Hadassah Hebrew University Medical Center and Faculty of Medicine, Jerusalem, Israel) for providing the *mTORC1* and *Tsc1* floxed mice and for valuable discussions and insight; Jie Li (Department of Genetics, University of Georgia, Athens, GA, USA, 30602, current address: NIAID, Bethesda, MD, USA, 20892) for technical assistance; E. Piontek, for technical assistance in paraffin embedding and tissue sectioning (Hadassah Hebrew University Medical Center, Israel). T. Naveh-Many is a research associate of the Wohl’s Translation Research Institute at Hadassah Hebrew University Medical Center.

## Supplemental Figure legends

**Supplemental Figure 1. Parathyroid-specific deletion of *Dicer* and miRNA in the parathyroid does not affect the expression of PTH or cleaved caspase3, nor the volume of the parathyroids at E19.5 and E21.5.** (A) Representative IF staining for PTH and cleaved caspase3 of parathyroids from control PT-*Dicer^+/+^* and PT-*Dicer^-/-^* embryos at both E19.5 (B) Similar IF staining images for parathyroids as in A at E21.5 (C&D) Quantitative volumetric assessment of parathyroids (PT-*Dicer^+/+^* n=4-7, PT-*Dicer^-/-^* n=5 parathyroids). The division into right and left glands is illustrated in panel D.

**Supplementary Figure 2. PT-*YFP*;*Dicer^-/-^* mice have normal basal serum PTH, calcium and phosphate levels but a muted increase in serum PTH after adenine rich high-phosphorus diet-induced CKD.** Control PT-*YFP;Dicer^+/+^* and PT-*YFP;Dicer^-/-^* mice were fed a control or an adenine-rich high phosphorus diet for 3 weeks to induce CKD and SHP. Serum levels of calcium (A), phosphate (B), urea nitrogen (BUN) (C) and PTH (D) were quantified. Data are presented as mean±SE. *, p<0.05 compared to PT-*YFP;Dicer^+/+^* control mice fed a control diet. #, p<0.05 compared to control mice with CKD.

**Supplemental Figure 3. Analysis of control probes in parathyroid glands by RNAscope.** Representative RNAscope analysis performed on parathyroid glands from 2 month-old control mice using positive and negative GCM2, Dicer and PTH probes and nuclear Dapi. n=2.

**Supplemental Figure 4. No change in PTH content in parathyroids of PT-*mTORC1^-/-^* mice compared to control PT-*mTORC1^-/-^* mice.** (**A**) Representative IF PTH staining of paraffin sections from both control and PT-*mTORC1^-/-^* mice. (**B**) Quantification of the findings depicted in A and additional sections obtained from additional mice. *, p<0.05.

**Supplemental Figure 5. Decreased PTH mRNA levels with unaltered CasR mRNA in PT-*mTORC1^-/-^* mice.** RNA extracted from thyroparathyroid tissue of 3 month-old control PT-*mTORC1^+/+^* and PT-*mTORC1^-/-^* mice was analyzed for PTH and CasR mRNA levels by qRT-PCR, normalized against β-actin. *, p<0.05.

**Supplemental Figure 6. Parathyroid-specific PT-*Tsc1^-/-^* mice have increased p-S6 (Ser235/236) levels in their parathyroids.** IF staining of parathyroid sections from control PT-*Tsc1^+/+^* and PT-*Tsc1^-/-^* mice highlighting pS6 (Ser235/236) (red) and nuclear sytox (green). (**B**) Quantitative assessment of pS6 staining intensity, measured using ImageJ software. *, p<0.05.

**Supplemental Figure 7. Visualization of parathyroid glands in a PT-*Tsc1^-/-^* mouse under florescence microscopy for TdTomato displayed without magnification.** The neck area of an 11 month PT-*Tsc1^-/-^* mouse depicted in Figure 5A, and a control PT-*Tsc1^+/+^* mouse at the same age, observed through fluorescence microscope for td-Tomato parathyroids at magnification x1.2 (left) and without magnification (right) demonstrating the size and intensity of the td-Tomato signal, which remains visible even without fluorescent light and no magnification.

**Supplemental Figure 8. Representative fluorescence imaging for parathyroid gland size estimation in PT-*Tsc1^+/+^* and PT-*Tsc1^-/-^* mice, as shown in** **Figure 5B**. Representative td-Tomato (left panels) and bright field images (right panels) used for estimating the size of parathyroid glands, outlined by broken lines and analyzed using ImageJ software. The left panels depict tdTomato florescence imaging at each age.

**Supplemental Figure 9. Increased PTH mRNA levels with unaltered CasR mRNA in PT-*Tsc1^-/-^* mice.** RNA was extracted from thyroparathyroid tissue of PT-*Tsc1^+/+^* and PT-*Tsc1^-/-^* mice at 3 months of age and analyzed for PTH and CasR mRNA levels by qRT-PCR, normalized for GAPDH. *, p<0.05.

