## Supplementary figures and images for "The essential roles of Dicer-mediated mTORC1 signaling in parathyroid gland integrity and function: Insights from genetic mouse models and clinical data"

### Supplemental Figure 1

Supplemental Figure 1

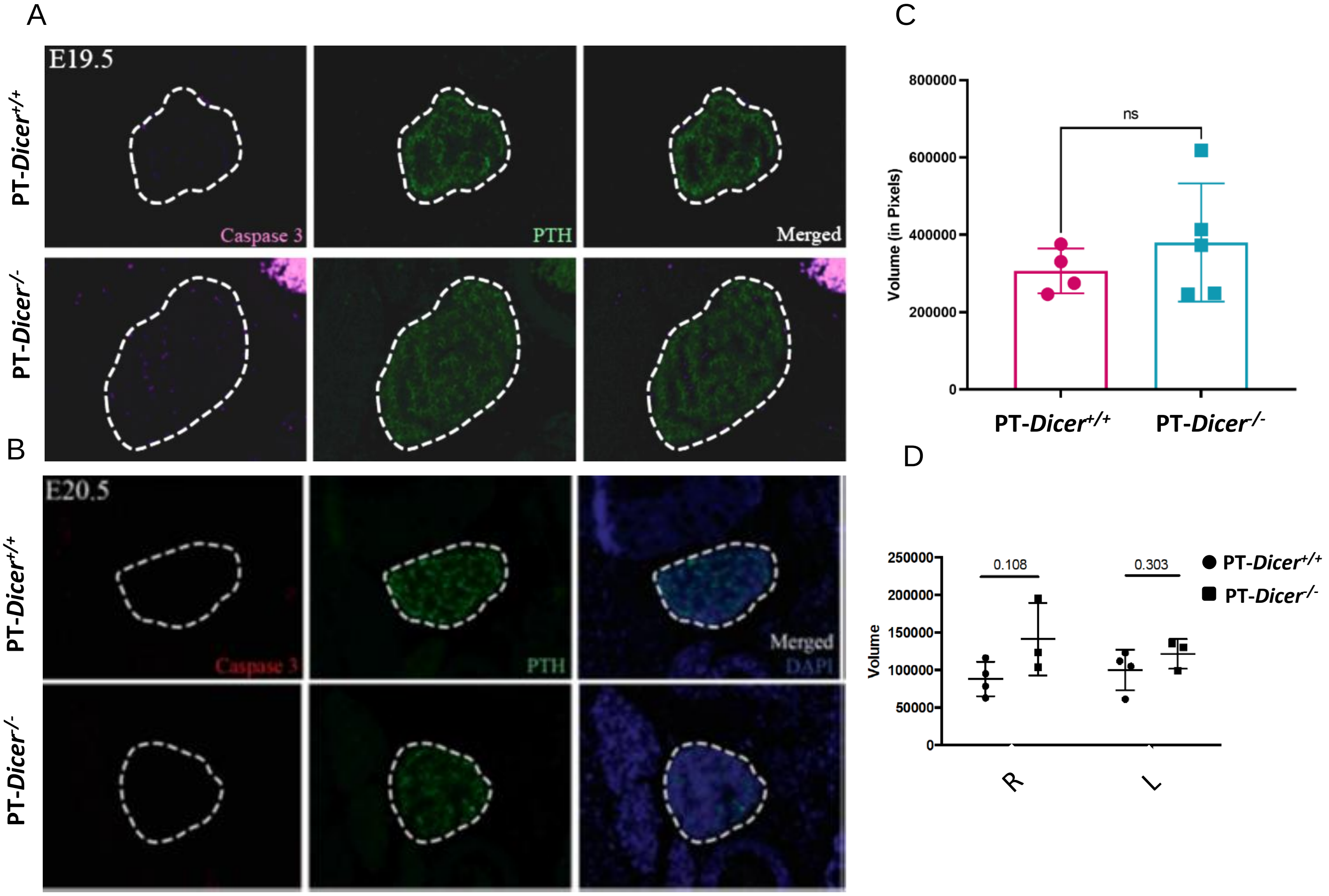

### Supplemental Figure 2

Supplemental Figure 2

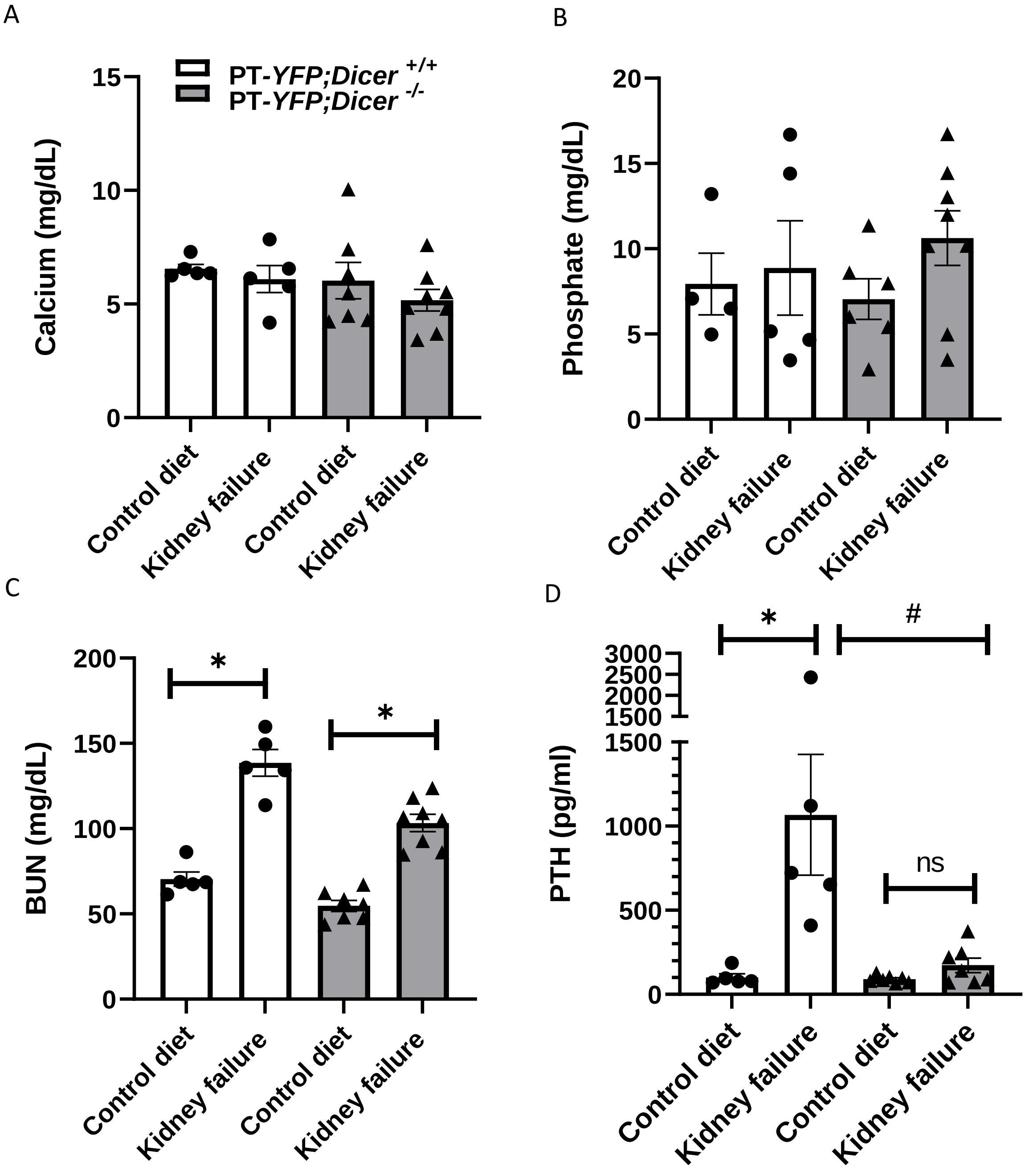

### Supplemental Figure 3

Supplemental Figure 3

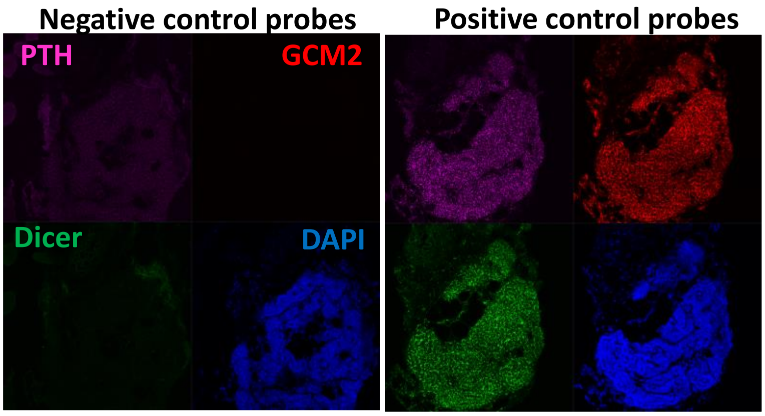

### Supplemental Figure 4

Supplemental Figure 4

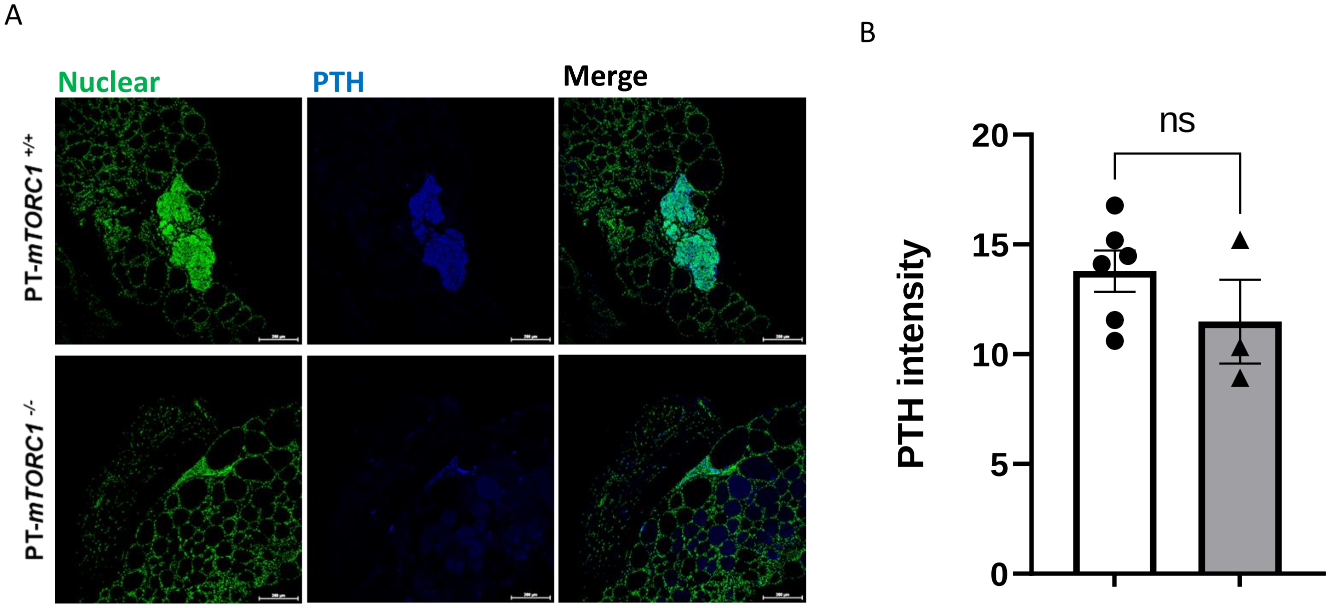

### Supplemental Figure 6

Supplemental Figure 6

A

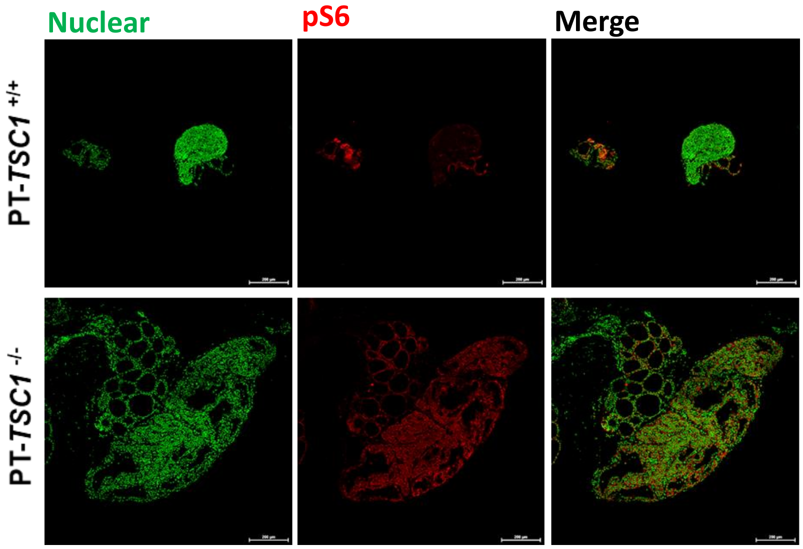


B

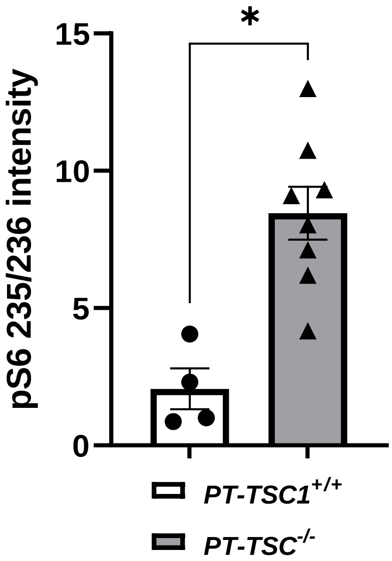

### Supplemental Figure 7

#### Supplemental Figure 7

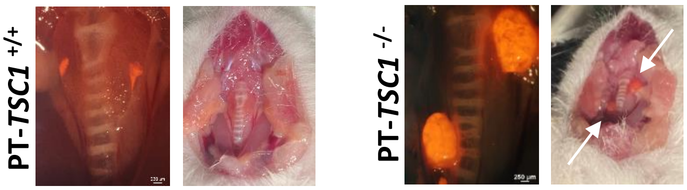

### Supplemental Figure 8

Supplemental Figure 8

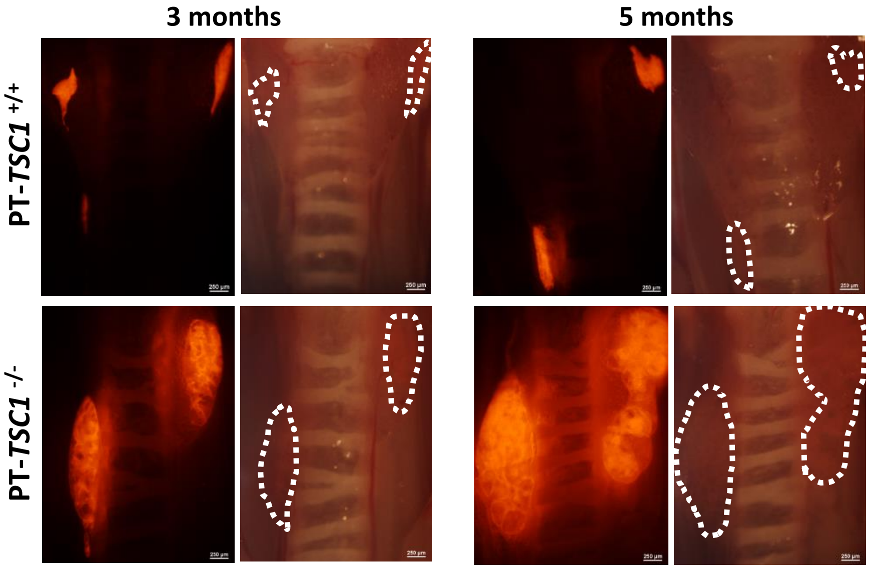

### Supplemental Figure 9

Supplemental Figure 9

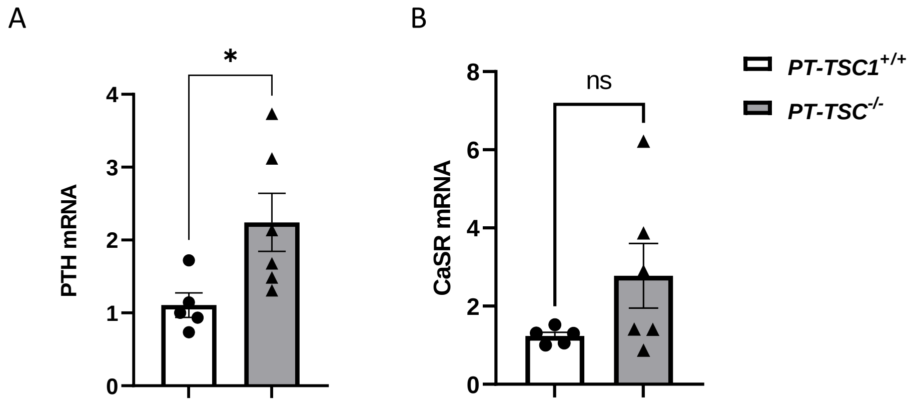
