## Supplemental Figure 5 for "The essential roles of Dicer-mediated mTORC1 signaling in parathyroid gland integrity and function: Insights from genetic mouse models and clinical data"

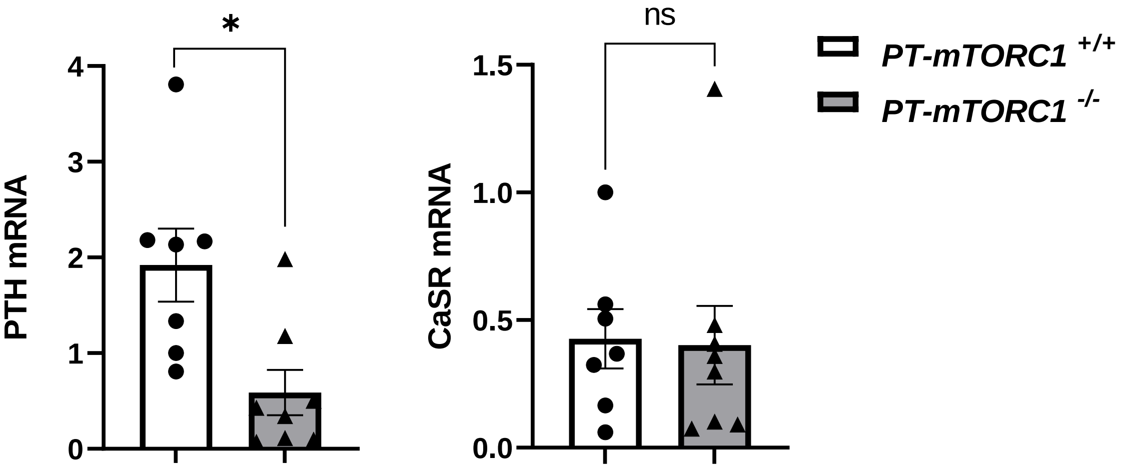


**Supplemental Figure 5. Decreased PTH mRNA levels with unaltered CasR mRNA in PT-*mTORC1*$^{-/-}$ mice.** RNA extracted from thyroparathyroid tissue of 3 month old control PT-*mTORC1*$^{+/+}$ and PT-*mTORC1*$^{-/-}$ mice was analyzed for PTH and CasR mRNA levels by qRT-PCR, normalized against β-actin. *, $p<0.05$.
