## Supplemental Mathods for "The essential roles of Dicer-mediated mTORC1 signaling in parathyroid gland integrity and function: Insights from genetic mouse models and clinical data"

### Supplemental Methods

#### Animal housing

All animal experiments were approved by the Institutional Animal Care and Use Committee of the Hadassah Hebrew University. Animals had free access to food and drinking water.

#### Parathyroid-specific knockout mice

Parathyroid-specific *Dicer* knockout mice (PT-*Dicer*$^{-/-}$) were generated by Cre-LoxP recombination using PTH-*Cre* mice, which express the *Cre* recombinase under the control of the human parathyroid hormone (PTH) promoter [FVBTg(PTH-Cre); Jackson Laboratory, Bar Harbor, ME] [38], as previously described [10]. To visualize the small mouse parathyroid glands using fluorescence microscopy, we introduced EYFP or tdTomato labeling. *EYFP* expression was achieved by crossing mice from both control PT-*Dicer*$^{+/+}$ and PT-*Dicer*$^{-/-}$ strains with *Rosa26 loxP*-STOP-*loxP EYFP* mutant mice (129X1-*Gt(ROSA)26Sor*$^{tm1(EYFP)Cos}$/J), from Jackson Laboratory, Bar Harbor, ME). This resulted in *Cre*-dependent *EYFP* expression specifically in the parathyroids (PT-*YFP;Dicer*$^{+/+}$ and PT-*YFP;Dicer*$^{-/-}$ mice). To enhance the detection of the small parathyroid cell clusters that were difficult to identify in YFP-expressing PT-*YFP;Dicer*$^{-/-}$ and PT-*mTORC1*$^{-/-}$ mice, we utilized the more intense tdTomato labeling. For this purpose, mice with a modified *Rosa26* locus17 were used, where the *Rosa26* locus contained a targeted insertion of a construct encompassing the strong and ubiquitously expressed *CAG* promoter, followed by a *loxP* stop cassette-controlled *tdTomato* marker gene [39]. Consequently, the resulting mice exhibited specific *tdTomato* expression in the parathyroids.

Parathyroid-specific *mTORC1* knockout mice (PT-*mTORC1*$^{-/-}$) and Parathyroid-specific *Tsc1* knockout mice (PT-*Tsc1*$^{-/-}$) were generated by crossing PTH-*Cre* mice with mice possessing floxed *mTORC1* [40, 41] *or Tsc1* genes [40, 42], respectively. The appropriate strains were further crossbred to generate the *Dicer* and *Tsc1* double knockout mice and their appropriate control mice (PT-*Dicer*$^{+/+}$;*Tsc1*$^{+/+}$ and PT-*Dicer*$^{-/-}$;*Tsc1*$^{-/-}$ mice). All control mice were littermates that were *Cre*-positive and had the appropriate wild type (wt) alleles for *Dicer*, *mTORC1* and *Tsc1*.

#### Genotyping

Total DNA was extracted from ear samples obtained from the offspring and genotyping was conducted using standard PCR techniques to identify the presence of

PTH-*Cre* and *Dicer loxP*, *YFP* and *tdTomato* alleles using the appropriate primers (Table 1) [10].

**Table 1. Primers used for DNA genotyping.**

| Genes | Sequence (5'-3') |
|---|---|
| PTH-*Cre* | Fw: TGCCACGACCAAGTGACAGC<br>Rev: CCAGGTTACGGATATAGTTCATG |
| *Dicer loxP* | Fw: CCTGACAGTGACGGTCCAAAG<br>Rev: CATGACTCTTCAACTCAAACT |
| *YFP* | ROSA 26R: AAAGTCGCTCTG AGTTGTTAT<br>BTG 60: GAAAGACCGCGAAGAGTT TG<br>BTG 62: TAAGCCTGCCCAGAAGACTC |
| *tdTomato* wt | Fw: AAGGGAGCTGCAGTGGAGTA<br>Rev: CCGAAAATCTGTGGGAAGTC |
| *tdTomato loxP* | Fw: CTGTTCCTGTACGGCATGG<br>Rev: GGCATTAAAGCAGCGTATCC |
| *mTOR loxP* | Fw: TTATGTTTGATAATTGCAGTTTTGGCTAG<br>Rev: TTAGGACTCCTTCTGTGACATACATTTC |
| *Tsc loxP* | Fw: AGGAGGCCTCTTCTGCTACC<br>Rev: CAGCTCCGACCATGAAGTG |

**Serum biochemistry and PTH levels**

Mice were anesthetized using a ketamine-xylazine solution, and blood samples were obtained from the abdominal aorta using heparin tubes. Following coagulation, the blood samples were centrifuged at 3000 g for 10 minutes to separate the serum, which was then isolated and stored at 20° C. Serum samples were analyzed using QuantiChrom kits (BioAssay Systems, Hayward, CA) to measure serum calcium and blood urea nitrogen (BUN) levels. In specific experiments, total serum calcium levels were determined using the Atellica CH Calcium kit (Tarrytown, NY). Serum PTH levels were measured using a mouse Intact 1–84 PTH ELISA kit (Quidel, Athens, OH) [10].

**Thyroparathyroidectomy**

The thyroparathyroidectomy procedure involved the removal of the thyroid gland along with the embedded parathyroid glands in control mice, or the parathyroid cell clusters as seen in PT-*Dicer*$^{-/-}$ mice. The mice, aged between 5 to 6 weeks, were anesthetized via intraperitoneal injection (IP) of ketamine-xylazine solution. A ventral midline incision was made in the neck area, and the brown adipose and muscle tissues were carefully retracted. Under a binocular microscope, the thyroids were located, exposed, and excised using fine scissors. Following the removal of the thyroid tissue, a sterile gauze was applied to the neck area, and the mice were kept warm under red light. For the sham-operated mice, the same procedure was performed, except the thyroids were spared. Thirty minutes after the thyroidectomy, blood samples were collected from both the thyroidectomized and sham-operated mice [20]. To confirm the successful removal of parathyroid-containing thyroid tissue in control mice, fluorescent-guided microscopy of the exposed neck was performed. Fluorescent-guided microscopy confirmed that PT-*YFP;Dicer*$^{-/-}$ mice lacked parathyroid glands altogether.

**RNA isolation and qRT-PCR**

Thyroparathyroid tissue was excised as above, rapidly frozen in liquid nitrogen and stored at -80° C to maintain its integrity. To extract total RNA from the two parathyroid glands of individual mice, 400 µl of TRIzol Reagent (Ambion, Carlsbad, CA) was utilized. The tissue was homogenized using a bead-beater, and RNA extraction was carried out in accordance with the manufacturer's instructions. The resulting RNA pellet was resuspended in 20 µl of RNase-free water, and 10 µl of the resuspended RNA was used for cDNA synthesis using a qScript cDNA Synthesis Kit (Quanta Bio, Beverly, MA). Subsequently, quantitative (q) PCR analysis was conducted using the PerfeCTa SYBR Green FastMix, Low ROX (Quanta Bio) in the ViiA 7 Fast Real-Time PCR System (Applied Biosystems, Waltham, MS). The primers used for qRT-PCR were for mouse PTH: Fw: 5'-ATCCCTTTGAGAGTCATTG-3' and Rev: 5'-GTTTGGGTAAGAAGACAGAC-3'; mouse calcium sensing receptor: Fw: 5'-GAGTGCATCAGGTATAACTTC-3' and Rev: 5'- GTGTTACAGGTGTCGAATATC-3' and mouse β-actin: Fw: 5'-CCTAGGCACCAGGGTGTGAT-3' and Rev: 5'-TCAGGGTCAGGATACCTCTCTTG-3' [10].

**Microscope fluorescent imaging and quantification**

Postnatal mice at various ages, ranging from P1 to adult, were anesthetized, and under bright field illumination, a surgical incision was made in the neck area to expose the thyroparathyroid. The parathyroid glands of PT-*YFP* or PT-*tdTomato* mice were visualized using GFP or CY3 filters, respectively, on a Nikon SMZ25 stereomicroscope. The exposure time for YFP fluorescence was set at 2 seconds, while for tdTomato, it was either 67 or 250 milliseconds as indicated, depending on the specific experiment requirements. The microscope images presented are composite images where the GFP or CY3 fluorescent signals were overlaid on bright field (BF) light images. To quantify the intensity of the fluorescent parathyroid gland images, ImageJ software (NIH, Bethesda, MD) was employed.

**Immunofluorescence staining of mouse paraffin sections**

Mouse thyroparathyroid glands were excised and fixed in 4% formaldehyde solution overnight. Subsequently the fixed glands were embedded in paraffin, and 6 µm-thick sections were prepared for further analysis. After rehydration, antigens were retrieved by pressure cooker (121° C for 3 min) in 20 mM citric acid buffer (pH= 6). Immunostaining was performed overnight at 4° C using primary antibodies for PTH (1:400, Bio-Rad, Hercules, CA), Cleaved caspase 3 (1:200, Cell Signaling Technology Danvers, MA), GCM2 (1:1000, Abcam, Cambridge, UK) CaSR (1:1500, Novus Biologicals, Littleton, CO), and phosphorylated ribosomal protein S6 (pS6) (S235/236) (1:500, Cell Signaling Technology) all diluted in Cas block (Zymed Laboratories, San Francisco, CA). Fluor chrome–conjugated secondary antibodies Cy3 and Cy5 (Bethyl, Montgomery, TX) and SYTOX nuclear staining (Life Technologies, Carlsbad, CA) were used for detection. Images were obtained using a Fluoview 1000 Olympus florescence microscope.

**Parathyroid gland size and *tdTomato* signal intensity**

The size of the parathyroid glands and the intensity of the fluorescent images were measured using ImageJ software (NIH, Bethesda, MD). 2D size estimation was performed on bright field images to avoid effects of the high intensity of tdTomato florescence (Supplemental Fig 8).

**Immunofluorescence staining of mouse embryos**

Embryos were obtained from timed matings at different stages of fetal parathyroid development. The day when a vaginal plug was observed was designated as embryonic day (E) 0.5. Embryos were collected on specific days of interest (E13.5, E19.5, and E21.5) and their developmental stage was determined based on somite counts and morphological features. For IF staining, 8 µm-thick serial sections of paraffin-embedded embryos, which were fixed in 4% paraformaldehyde (PFA), were used. The slides were subjected to staining for PTH (Quidel), CC3 (Cell Signaling Technology), and nuclear staining with DAPI.

Embryos for paraffin sections to be used in RNAscope analysis were fixed in a 4% paraformaldehyde solution overnight. Subsequently, they were thoroughly washed with PBS, dehydrated in a series of ethanol dilutions, permeabilized using xylene, and finally embryos were embedded in paraffin wax. All steps were performed using RNase-free reagents. Sections for RNAscope analysis were sliced at 5 µm. IHC was performed using antibodies for PTH (Quidel 21-2310), caspase 3 (Cell Signaling Technologies 9661S), both of which were diluted at 1:200 and DAPI nuclear staining.

Using IHC-generated images, three-dimensional (3D) reconstructions of the parathyroids were created utilizing the WinSurf® software. These reconstructions enabled the generation of volumetric data for all collected parathyroids. The embryos were examined at three developmental stages: E13.5, E19.5, and E21.5. To quantify the size of the parathyroid domain, the sum of all areas (in pixels) within a pouch was calculated. These calculations were then entered into Prism© software for subsequent statistical analysis. The statistical significance was determined using an unpaired T-test. For the E13.5, 8 control embryos (16 parathyroids) and 15 mutant embryos (30 parathyroids) were examined using IHC. Among these, 4 control parathyroids and 7 mutant parathyroids were quantified. Similarly, for the E19.5 stage, 4 control embryos (8 parathyroids) and 7 mutant embryos (14 parathyroids) were subjected to IHC analysis. Quantification was performed on 4 control parathyroids and 5 mutant parathyroids. At the E21.5 stage, 5 control embryos (10 parathyroids) and 6 mutant embryos (12 parathyroids) were examined.

**RNAscope in situ hybridization**

RNAscope analysis was conducted on RNase-free paraformaldehyde-fixed, paraffin-embedded sections following the protocols and utilizing the reagents provided by the manufacturer RNAscope™ (ACDbio, Newark, CA) [43]. The specific probes used included Gcm2, Pth and Dicer.

**Human clinical investigation**

To explore the clinical relationship between mTOR inhibition and serum PTH levels, we conducted an analysis involving human kidney transplant recipients. Our study compared patients who were undergoing immunosuppressive treatment regimens that included mTOR inhibitors with those receiving calcineurin inhibitors. We accessed summary data from a substantial population of kidney transplant patients through the TriNetX network datasets [44]. We collected data as of 5/Sep/2023 from the TriNetX Global Collaborative Network This network provides access to statistics on comprehensive array of electronic medical records, encompassing diagnoses, procedures, medications, laboratory values, and genomic information, from a vast patient pool, comprising over 127 million individuals, representing data derived from 106 healthcare organizations. As a federated network [44], TriNetX received a waiver from Western IRB since only aggregated counts, statistical summaries of de-identified information, but no protected health information is received, and no study-specific activities are performed in retrospective analyses.

**Statistical analysis**

Values are presented as mean±SE. A 2-tailed Student's t-test was used to assess differences among groups. A p value of less than 0.05 was considered significant.
